## Supplementary material for "A divide-and-conquer method for analyzing high-dimensional noisy gene expression networks": SI Appendix

#### This PDF file includes:

Tables S1 to S2 (not allowed for Brief Reports)
SI References

**S8 Noise decomposition based on ergodicity** **25**

 Zhou Fang, Ankit Gupta, Sant Kumar, Mustafa Khammash

### S1. Stochastic intracellular reaction systems, the chemical master equation (CME), and the associated stochastic filtering problem

In this work, we focus on an intracellular reaction system that has $r$ reactions:

$$v_{1,j}S_1 + \cdots + v_{n,j}S_n \to v'_{1,j}S_1 + \cdots + v'_{n,j}S_n \qquad j = 1, \ldots, r, \qquad [1]$$

where $S_1, \ldots, S_n$ are $n$ different chemical species, and $v_{i,j}$ and $v'_{i,j}$ are the stoichiometric coefficients indicating the number of molecules consumed or produced for $S_i$ in the $j$-th reaction. Due to the low molecular counts, an intracellular reaction system is inevitably random, and, therefore, its dynamics is usually modeled by a stochastic process called the continuous time Markov chain. Following (1), the dynamical equation can be written by

$$X(t) = X(0) + \sum_{j=1}^{r} \zeta_j R_j \left( \int_0^t \lambda_j(X(s))\mathrm{d}s \right) \qquad [2]$$

where $X(t)$ is an $n$-dimensional vector representing the molecular count for each species at time $t$, the vector $\zeta_j$ equals to $\left(v'_{1,j} - v_{1,j}, \ldots, v'_{1,n} - v_{1,n}\right)^\top$ indicating the state change after a firing of the $j$-th reaction, $R_j(t)$ are independent unit rate Poisson processes, and $\lambda_j(\cdot)$ are the propensities indicating the rates of these reactions. To avoid negative molecular counts, the propensities should satisfy the condition $\lambda_j(x) = 0$ for all $x + \zeta_j \notin \mathbb{Z}^n_{\geq 0}$. For rigorousness, we only consider the processes that satisfy the condition

$$\sum_{j=1}^{n} \mathbb{E}\left[\lambda_j^2(X(t))\right] \text{ is uniformly bounded on any time interval } [0, T], \qquad [3]$$

which means the system state will almost surely not grow to infinity in finite time. What's more, this condition also implies that the random variables $\{\lambda_j(X(t))\}_{t\in[0,T]}$ are uniformly integrable. Also, for simplicity, we assume that the initial conditions for different species are independent, i.e.,

$$X_1(0), \ldots, X_n(0) \text{ are independent.} \qquad [4]$$

Alternative to Eq. [2], one can also model the intracellular reaction system by the chemical master equation (CME) (1):

$$\frac{\mathrm{d}p(t,x)}{\mathrm{d}t} = \sum_{j=1}^{r} \lambda_j(x - \zeta_j)p(t, x - \zeta_j) - \sum_{j=1}^{r} \lambda_j(x)p(t,x), \qquad \forall x \in \mathbb{Z}_{\geq 0},$$

where $p(t,x) \triangleq \mathbb{P}(X(t) = x)$. Provided with the condition Eq. [3], these two dynamical presentations are equivalent (1).

In the lab, scientists can directly measure fluorescent reporters and use these measurements to infer the dynamical states of unobserved species. This is called the stochastic filtering for intracellular reaction systems. Mathematically, we can model the observations by

$$Y(t_i) = h(X(t_i)) + \sigma W_i \qquad [5]$$

where $t_i$ are the observation time points, $Y(t_i)$ is a vector of observations with each element corresponding to a particular light frequency, $h(\cdot)$ is a measurable function, $\sigma$ is a diagonal matrix indicating observation noise intensities, and $W_i$ are independent Gaussian white noise. The goal of the stochastic filtering is to compute the conditional expectation $\pi_{t_i}(x) \triangleq \mathbb{P}\left(X(t_i) = x \middle| Y(t_s),\ 1 \leq s \leq i\right)$. By the Bayes' rule, the solution of the filtering problem satisfies the following recursive formulas (2)

$$\rho_{t_{i+1}}(x) = \sum_{x' \in \mathbb{Z}^n_{\geq 0}} \mathbb{P}\left(X(t_{i+1}) = x \middle| X(t_i) = x'\right) \pi_{t_i}(x') \qquad [6]$$

$$\pi_{t_{i+1}}(x) \propto L\left(Y(t_{i+1}) \middle| x\right) \rho_{t_{i+1}}(x) \qquad [7]$$

where $\rho_{t_{i+1}}(x) \triangleq \mathbb{P}\left(X(t_{i+1}) = x \middle| Y(t_s),\ 1 \leq s \leq i\right)$ and $L\left(y|x\right)$ is the density function of $\mathbb{P}\left(Y(t_{i+1}) \in \cdot \middle| X(t_{i+1}) = x\right)$ (usually called the likelihood function). We can interpret Eq. [6] as the prediction of the state at the next time point $t_{i+1}$ using the observation up to the current time $t_i$, and we interpret Eq. [7] as the adjustment of the predication according to the new observations. Note that the prediction step Eq. [6] is actually solving a CME with $\pi_{t_i}(\cdot)$ being the initial probability and $\rho_{t_{i+1}}(\cdot)$ being the final solution. Consequently, the filtering problem can be seen as a combination of the CME and an adjustment step.

### S2. Error analysis of the Monte-Carlo method and the particle filter

One popular method to solve the CME is the Monte-Carlo method, which utilizes stochastic simulations to approximate the exact probability. Since the stochastic filtering problem can be seen as a combination of the CME and an adjustment step, one can also solve the filtering problem by using the Monte Carlo for the CME step. In the literature, this Monte-Carlo method for the filtering problem is called the particle filter or the sequential Monte-Carlo method.

In this section, we perform the error analysis of these Monte-Carlo methods and show that their error tends to grow exponentially with the system dimension. Moreover, we also show that both approaches have relatively the same performance when solving their associated problems.

**A. Error analysis of the Monte-Carlo method for CMEs.** To solve a CME, the Monte-Carlo method first simulates $N$ trajectories of the system Eq. [2], denoted by $\mathrm{x}_1(t), \dots, \mathrm{x}_N(t)$, and then uses the empirical distribution $p_{\mathrm{MC}}(t,x) \triangleq \sum_{j=1}^{N} \mathbb{1}\left(\mathrm{x}_1(t) = x\right)$ to approximate the exact probability. Here, $\mathbb{1}(\cdot)$ is the indicator function whose value equals to 1 if its argument is true, otherwise 0. Also, for simplicity, we only use the exact simulation method (e.g., Gillespie method) to generate the trajectories.

We now analyze the $L_1$ error of the Monte-Carlo method, defined by $\left\|\hat{p}_{\mathrm{MC}}(t,\cdot) - p(t,\cdot)\right\|_1 \triangleq \sum_{x \in \mathbb{Z}^n_{\geq 0}} \left|\hat{p}_{\mathrm{MC}}(t,x) - p(t,x)\right|$. By the law of the central limit theorem, we have that for each $x \in \mathbb{Z}^n_{\geq 0}$,

$$\sqrt{N}\left(\hat{p}_{\mathrm{MC}}(t,x) - p(t,x)\right) \xrightarrow{d} \mathcal{N}\left(0, \mathrm{Var}\left(\mathbb{1}\left(\mathrm{x}_1(t) = x\right)\right)\right), \quad \text{as } N \to \infty.$$

Notice that $\mathbb{1}\left(\mathrm{x}_1(t) = x\right)$ has the Bernoulli distribution, so $\mathrm{Var}\left(\mathbb{1}\left(\mathrm{x}_1(t) = x\right)\right) = p(t,x)\left(1 - p(t,x)\right)$ for every $x \in \mathbb{Z}^n_{\geq 0}$. Therefore, if the sum $\sum_{x \in \mathbb{Z}^n_{\geq 0}} \sqrt{p(t,x)}$ is convergent, we have that

$$\lim_{N\to\infty} \sqrt{N}\mathbb{E}\left[\left\|\hat{p}_{\mathrm{MC}}(t,\cdot) - p(t,\cdot)\right\|_1\right] = \lim_{N\to\infty} \sqrt{N} \sum_{x \in \mathbb{Z}^n_{\geq 0}} \mathbb{E}\left[\left|\hat{p}_{\mathrm{MC}}(t,x) - p(t,x)\right|\right] = \sqrt{\frac{2}{\pi}} \sum_{x \in \mathbb{Z}^n_{\geq 0}} \sqrt{p(t,x)(1 - p(t,x)}. \qquad [8]$$

Finally, by the relation $\sqrt{a-b} \geq \sqrt{a} - \sqrt{b}$ for any $0 \leq b \leq a$, we can conclude

$$\lim_{N\to\infty} \sqrt{N}\mathbb{E}\left[\left\|\hat{p}_{\mathrm{MC}}(t,\cdot) - p(t,\cdot)\right\|_1\right] = \sqrt{\frac{2}{\pi}}\left[\left(\sum_{x \in \mathbb{Z}^n_{\geq 0}} \sqrt{p(t,x)}\right) \pm 1\right], \qquad [9]$$

which provides upper and lower bounds for the error.

The formula Eq. [9] tells that when the sample size $N$ is large and fixed, the error of the Monte-Carlo method largely depends on the value of $\sum_{x \in \mathbb{Z}^n_{\geq 0}} \sqrt{p(t,x)}$. This value tends to scale poorly with the size of the state space containing most of the probability mass. Particularly, when the probability mass is uniformly distributed on $T$ states, this quantity equals $\sqrt{T}$, which can be very large when $T$ is big. Also, since this state size often grows exponentially with the number of species, this quantity $\sum_{x \in \mathbb{Z}^n_{\geq 0}} \sqrt{p(t,x)}$ tends to scale poorly with the system dimension $n$. When the molecular counts of different chemical species are independent, this sum does grow exponentially with $n$, as its value equals to $\prod_{i=1}^{n}\left(\sum_{x_i \in \mathbb{Z}_{\geq 0}} \sqrt{p_i(t,x_i)}\right)$ with $p_i(t,\cdot)$ the marginal distribution for the $i$-th species. In summary, the error of a Monte-Carlo method tends to scale unfavorably with the size of state space and system dimension, and, thus, this method usually performs poorly in high-dimensional problems.

Next, we show that if the state variables are independent and their marginal distributions are estimated separately, then the errors in estimating these marginal distributions additively contribute to the error in the joint probability estimate. Again, let us denote $p_i(t,x_i)$ the marginal distribution for the $i$-th species, and $\hat{p}_i(t,x)$ be an estimate of this marginal distribution by the Monte-Carlo method. Here, we consider that these marginal distributions are estimated separately, using different sets of

 Zhou Fang, Ankit Gupta, Sant Kumar, Mustafa Khammash

samples. Therefore, these estimates are probabilistically independent, and we can get

$$
\begin{aligned}
&\underbrace{\sum_{x_1\in\mathbb{Z}_{\geq 0}}\cdots\sum_{x_n\in\mathbb{Z}_{\geq 0}}\mathbb{E}\left[\left|\prod_{i=1}^{n}\hat{p}_i(t,x_i)-\prod_{i=1}^{n}p_i(t,x_i)\right|\right]}_{\text{Error in estimating the joint distribution}}\\
&\leq \sum_{x_1\in\mathbb{Z}_{\geq 0}}\cdots\sum_{x_n\in\mathbb{Z}_{\geq 0}}\mathbb{E}\left[\hat{p}_1(t,x_1)\left|\prod_{i=2}^{n}\hat{p}_i(t,x_i)-\prod_{i=2}^{n}p_i(t,x_i)\right|+\left|\hat{p}_1(t,x_1)-p_1(t,x_1)\right|\prod_{i=2}^{n}p_i(t,x_i)\right]\\
&=\sum_{x_2\in\mathbb{Z}_{\geq 0}}\cdots\sum_{x_n\in\mathbb{Z}_{\geq 0}}\mathbb{E}\left[\left|\prod_{i=2}^{n}\hat{p}_i(t,x_i)-\prod_{i=2}^{n}p_i(t,x_i)\right|\right]+\sum_{x_1\in\mathbb{Z}_{\geq 0}}\mathbb{E}\left[\left|\hat{p}_1(t,x_1)-p_1(t,x_1)\right|\right] \qquad \text{(by independence)}\\
&\ldots\ldots\\
&\leq \sum_{i=1}^{n}\underbrace{\left[\sum_{x_i\in\mathbb{Z}_{\geq 0}}\mathbb{E}\left[\left|\hat{p}_i(t,x_i)-p_i(t,x_i)\right|\right]\right]}_{\text{Error in estimating each marginal distribution}}.
\end{aligned}
$$

This result suggests that the error in estimating the joint distribution is no greater than the sum of the errors in estimating marginal distributions.

**B. Error analysis of the particle filter.** Recall that the filtering problem can be seen as a combination of the CME and the adjustment step. Therefore, one idea to solve this problem is using the Monte-Carlo method for the CME step. In the literature, this method is called the particle filter, whose detailed algorithm is summarized in Algorithm 1. Specifically, the particle filter first generates equally weighted particles $(\mathrm{x}_1(0),\ldots \mathrm{x}_N(0))$ from the initial probability so that their empirical distribution approximates the initial distribution. Then, in each iteration, the algorithm simulates the particles to solve the CME in the prediction step and then correct their weights according to the new observations. Based on the prediction and the correction steps, a numerical filter is constructed by the empirical distribution of the particles, i.e., $\hat{\pi}^{\mathrm{PF}}_{t_{i+1}}(x)=\sum_{j=1}^{N}w_j(t_{i+1})\mathbb{1}\left(\mathrm{x}_j(t_{i+1})=x\right)$. Finally, the algorithm resamples particles to delete non-significant particles.

---

**Algorithm 1** Particle filter (adapted from (3))

---

Generate $N$ particle $(\mathrm{x}_1(0),\ldots,\mathrm{x}_N(0))$ from the initial probability, and give them equal weights ($w_j(0)=\frac{1}{N}$, $j=1,\ldots,N$). ▷ Initialization
$i\leftarrow 0$ and $t_0\leftarrow 0$.
**while** $t_i$ is not the final observation time **do**
  Simulate every particle from time $t_i$ to $t_{i+1}$ according to Eq. [2]. ▷ Prediction (see Eq. [6])
  Update weights $w_j(t_{i+1})\propto w_j(t_i)L(Y(t_{i+1})|\mathrm{x}_j(t_{i+1}))$. ▷ Adjustment (see Eq. [7])
  Compute the filter $\hat{\pi}^{\mathrm{PF}}_{t_{i+1}}(x)=\sum_{j=1}^{N}w_j(t_{i+1})\mathbb{1}\left(\mathrm{x}_j(t_{i+1})=x\right)$ ▷ Compute the filter
  Resample $\{w_j(t_{i+1}),x_j(t_{i+1})\}$ to obtain equally weighted particles $\{1/N,x_j(t_{i+1})\}$ ▷ Delete non-significant particles
  $i\leftarrow i+1$.
**end while**

---

Now, we analyze the error of the particle filter $\hat{\pi}^{\mathrm{PF}}_{t_i}(x)$. We first look at the $L_1$ error of this filter at the first observation time point $t_1$. (Readers who are not interested in the mathematical derivation can directly go to Eq. [13].) Given a fixed $Y(t_i)$, the literature (4, Theorem 1) tells that for any $x\in\mathbb{Z}^n$, there is the central limit result

$$
\sqrt{N}\left(\hat{\pi}^{\mathrm{PF}}_{t_1}(x)-\pi_{t_1}(x)\right)\xrightarrow{\text{in distribution}}\mathcal{N}\left(0,\mathbb{V}ar\left[\frac{\pi_{t_1}\left(\mathrm{x}_1(t_1)\right)}{\rho_{t_1}\left(\mathrm{x}_1(t_1)\right)}\left[\mathbb{1}\left(\mathrm{x}_1(t_1)=x\right)-\pi_{t_1}(x)\right]\middle|Y(t_1)\right]\right), \qquad \text{as } N\to\infty,
$$

where $\pi_{t_1}(\cdot)$ is the exact filter at $t_1$, the letter $\mathcal{N}$ is the notation for the Gaussian distribution, the term $\mathrm{x}_1(t_1)$ is the value of the first particle at time $t_1$ before the resampling step, the term $\rho_{t_1}(x)$ is the prediction probability $\mathbb{P}\left(X(t_1)=x\right)$, and $\mathbb{1}(\cdot)$ is the indicator function. Based on this formula, we can conclude that

$$
\lim_{N\to\infty}\sqrt{N}\,\mathbb{E}\left[\left|\hat{\pi}^{\mathrm{PF}}_{t_1}(x)-\pi_{t_1}(x)\right|\middle|Y(t_1)\right]=\sqrt{\frac{2}{\pi}}\sqrt{\mathbb{V}ar\left[\frac{\pi_{t_1}\left(\mathrm{x}_1(t_1)\right)}{\rho_{t_1}\left(\mathrm{x}_1(t_1)\right)}\left[\mathbb{1}\left(\mathrm{x}_1(t_1)=x\right)-\pi_{t_1}(x)\right]\middle|Y(t_1)\right]}. \qquad [10]
$$

To further investigate Eq. [10], we simplifies the variance in this formula as follows. We first term $\rho^{Y}_{t_1}(y)\triangleq\sum_{x\in\mathbb{R}^n}L(y|x)\rho_{t_1}(x)$ as the density of $Y(t_1)$. Using this notation, we can write $\pi_{t_1}\left(\mathrm{x}_1(t_1)\right)/\rho_{t_1}\left(\mathrm{x}_1(t_1)\right)=L\left(Y(t_1)\middle|\mathrm{x}_1(t_1)\right)/\rho^{Y}_{t_1}\left(Y(t_1)\right)$. Since $\mathrm{x}_1(t_1)$

has the distribution $\rho_{t_1}$, we can easily check that given $Y(t_1)$, the random variable $\frac{\pi_{t_1}(\mathrm{x}_1(t_1))}{\rho_{t_1}(\mathrm{x}_1(t_1))}\left[\mathbb{1}\left(\mathrm{x}_1(t_1)=x\right)-\pi_{t_1}(x)\right]$ has zero conditional mean, and its conditional variance satisfies

$$\mathbb{V}ar\left[\frac{\pi_{t_1}\left(\mathrm{x}_1(t_1)\right)}{\rho_{t_1}\left(\mathrm{x}_1(t_1)\right)}\left[\mathbb{1}\left(\mathrm{x}_1(t_1)=x\right)-\pi_{t_1}(x)\right]\middle|Y(t_1)\right]\leq\left(\frac{L\left(Y(t_1)\middle|x\right)}{\rho^Y_{t_1}\left(Y(t_1)\right)}\right)^2\rho_{t_1}(x)+\pi^2_{t_1}(x)\sum_{x'\in\mathbb{R}^n}\left(\frac{L\left(Y(t_1)\middle|x'\right)}{\rho^Y_{t_1}\left(Y(t_1)\right)}\right)\pi_{t_1}(x')$$

and

$$\mathbb{V}ar\left[\frac{\pi_{t_1}\left(\mathrm{x}_1(t_1)\right)}{\rho_{t_1}\left(\mathrm{x}_1(t_1)\right)}\left[\mathbb{1}\left(\mathrm{x}_1(t_1)=x\right)-\pi_{t_1}(x)\right]\middle|Y(t_1)\right]\geq\left(\frac{L\left(Y(t_1)\middle|x\right)}{\rho^Y_{t_1}\left(Y(t_1)\right)}\right)^2\rho_{t_1}(x)\left(1-\pi_{t_1}(x)\right)^2.$$

Moreover, by applying $\sqrt{a+b}\leq\sqrt{a}+\sqrt{b}$ and $L(y|x)\leq\det(\sqrt{2\pi\sigma})^{-1/2}$ (det is the determinant notation) to the above equations, we can further obtain

$$\sqrt{\mathbb{V}ar\left[\frac{\pi_{t_1}\left(\mathrm{x}_1(t_1)\right)}{\rho_{t_1}\left(\mathrm{x}_1(t_1)\right)}\left[\mathbb{1}\left(\mathrm{x}_1(t_1)=x\right)-\pi_{t_1}(x)\right]\middle|Y(t_1)\right]}\leq\left(\frac{L\left(Y(t_1)\middle|x\right)}{\rho^Y_{t_1}\left(Y(t_1)\right)}\right)\sqrt{\rho_{t_1}(x)}+\pi_{t_1}(x)\sqrt{\frac{\det(\sqrt{2\pi\sigma})^{-1/2}}{\rho^Y_{t_1}\left(Y(t_1)\right)}}\qquad[11]$$

and

$$\begin{aligned}\sqrt{\mathbb{V}ar\left[\frac{\pi_{t_1}\left(\mathrm{x}_1(t_1)\right)}{\rho_{t_1}\left(\mathrm{x}_1(t_1)\right)}\left[\mathbb{1}\left(\mathrm{x}_1(t_1)=x\right)-\pi_{t_1}(x)\right]\middle|Y(t_1)\right]}&\geq\left(\frac{L\left(Y(t_1)\middle|x\right)}{\rho^Y_{t_1}\left(Y(t_1)\right)}\right)\sqrt{\rho_{t_1}(x)}\left(1-\pi_{t_1}(x)\right)\\&=\left(\frac{L\left(Y(t_1)\middle|x\right)}{\rho^Y_{t_1}\left(Y(t_1)\right)}\right)\sqrt{\rho_{t_1}(x)}\left(1-\sqrt{\rho_{t_1}(x)}\sqrt{\pi_{t_i}(x)}\sqrt{\frac{L\left(Y(t_1)\middle|x\right)}{\rho^Y_{t_1}\left(Y(t_1)\right)}}\right)\\&\geq\left(\frac{L\left(Y(t_1)\middle|x\right)}{\rho^Y_{t_1}\left(Y(t_1)\right)}\right)\left(\sqrt{\rho_{t_1}(x)}-\rho_{t_1}(x)\sqrt{\frac{\det(\sqrt{2\pi\sigma})^{-1/2}}{\rho^Y_{t_1}\left(Y(t_1)\right)}}\right).\qquad[12]\end{aligned}$$

Finally, by plugging Eq. [11] and Eq. [12] into Eq. [10], we can estimate the $L_1$ error of the particle filter by

$$\begin{aligned}\lim_{N\to\infty}\sqrt{N}\,\mathbb{E}\left[\left\|\hat{\pi}^{\mathrm{PF}}_{t_1}-\pi_{t_1}\right\|_1\right]&\triangleq\lim_{N\to\infty}\sqrt{N}\sum_{x\in\mathbb{Z}^n}\mathbb{E}\left[\left|\hat{\pi}^{\mathrm{PF}}_{t_1}(x)-\pi_{t_1}(x)\right|\right]\\&=\sqrt{\frac{2}{\pi}}\left[\left(\sum_{x\in\mathbb{Z}^n_{\geq0}}\sqrt{\rho_{t_1}(x)}\right)\pm\det(\sqrt{2\pi\sigma})^{-1/4}\int_{y\in\mathbb{R}^m}\sqrt{\rho^Y_{t_1}(y)}\,\mathrm{d}y\right],\qquad[13]\end{aligned}$$

provided that the quantities $\sum_{x\in\mathbb{Z}^n_{\geq0}}\sqrt{\rho_{t_1}(x)}$ and $\int_{y\in\mathbb{R}^m}\sqrt{\rho^Y_{t_1}(y)}\,\mathrm{d}y$ are convergent. Here, $m$ is the dimension of $Y(t_1)$.

The formula Eq. [13] tells that for a large but fixed sample size $N$, the error of the particle filter at time $t_1$ largely depends on the quantities $\sum_{x\in\mathbb{Z}^n_{\geq0}}\sqrt{\rho_{t_1}(x)}$ and $\det(\sqrt{2\pi\sigma})^{-1/4}\int_{y\in\mathbb{R}^m}\sqrt{\rho^Y_{t_1}(y)}\,\mathrm{d}y$. Similar to the analysis in the previous subsection, these two terms tend to grow exponentially with the system dimension and the observation dimension, respectively. Note that the system dimension is usually much larger than the observation dimension, and the observation noise is usually not negligible. So, we can conclude that the quantity $\sum_{x\in\mathbb{Z}^n_{\geq0}}\sqrt{\rho_{t_1}(x)}$ dominates in Eq. [13], and, therefore, the $L_1$ error of $\pi^{\mathrm{PF}}_{t_1}$ tend to scale exponentially with the system dimension $n$. Moreover, this also suggests that the particle filter for the filtering problem has relatively the same performance as the Monte-Carlo method for solving the CME. Particularly, their performance are the same when the observation noise intensity is sufficiently large, i.e., the observation does not provide any information of the system.

**Remark 1.** *For other observation time points, we conjecture that the same results should also hold true. A theoretical analysis can be performed in the same way as we presented above, with the exception that the particles* $\mathrm{x}_1(t_i),\ldots,\mathrm{x}_N(t_i)$ *do not have the distribution* $\rho_{t_i}$ *before the adjustment step. Instead, the particles converges to the distribution* $\rho_{t_i}$ *in the limit of large sample size (4), which definitely complicates the analysis. However, we conjecture that this discrepancy should not affect the performance of the particle filer too much, and, therefore, the results obtained for time* $t_1$ *should also apply to other time points. Our numerical studies also support this point (see Section 2.C in the main text). Since this theoretical analysis can be very long, we leave it for further work.*

 

### S3. Rao-Blackwellized CME solver (RB-CME solver): derivation, error analysis, and the optimal lead-follower decomposition.

**A. Leader-follower decomposition and the conditional independence among follower subsystems.** The Rao-Blackwellized CME solver (RB-CME solver) is established based on a decomposition of the system. We first divide the system state $X(t)$ into a leader system $\tilde{X}(t)$ and a follower subsystem $Z(t)$. Furthermore, we decompose each reaction vector $\zeta_j$ into $\zeta_j^{\tilde{X}}$ and $\zeta_j^{Z}$, where $\zeta_j^{\tilde{X}}$ indicates the state change of the leader system, and $\zeta_j^{Z}$ indicates the state change of the follower system. For the follower system, we further decompose it into several subsystems, $Z_1(t), \ldots, Z_l(t)$, such that the following topological conditions are satisfied.

C1 Each reaction involves a maximum of one follower subsystem (meaning that at most one follower subsystem can influence the reaction's propensity or have its state altered by the reaction).

C2 The reactions with the same non-zero $\zeta_j^{\tilde{X}}$ involve a maximum of one follower subsystem (meaning that at most one follower subsystem can influence the propensities of these reactions or have its state altered by these reactions).

For the ease of notations, we rearrange the order of species such that $X(t) = \left(\tilde{X}(t), Z_1(t), \ldots, Z_l(t)\right)$; also, for every state $x$, we write $x = (\tilde{x}, z_1, \ldots, z_l)$, where $\tilde{x}$ is the state of the leader system, and $z_i$ $(i = 1, \ldots, l)$ is the state of the follower system. We denote the dimension of the leader system by $n_{\tilde{X}}$ and the dimension of the follower system by $n_Z$. An automated algorithm for this leader-follower decomposition is presented in Section S3.D.

One key ingredient in our new method is computing the conditional distribution of the follower system given the trajectory of the leader one, i.e, $\mathbb{P}\left(Z(t) = z \middle| \tilde{X}(s),\ 0 \le s \le t\right)$. Mathematically, this conditional probability is not uniquely defined for each $t > 0$; it can be any $\tilde{\mathcal{X}}_t$-measurable* random variable $\mathcal{P}(z)$ satisfying

$$\mathbb{E}\left[\mathcal{P}(z) A\right] = \mathbb{E}\left[\mathbb{1}(Z(t) = z) A\right] \qquad \forall \tilde{\mathcal{X}}_t\text{-measurable random variable } A \text{ with finite expectation.}$$

Therefore, $\mathbb{P}\left(Z(t) = z \middle| \tilde{X}(s),\ 0 \le s \le t\right)$ viewed as a continuous-time process is not uniquely defined either, and some of them can be non-cadlag. ("Cadlag" means right continuous and having a left limit). Fortunately, ([5], Theorem 2.24) and the right continuity of $X(t)$ guarantee that for each $z \in \mathbb{Z}_{\ge 0}^{n_Z}$, there exists a $\tilde{\mathcal{X}}_t$-adaptive cadlag process $\pi_{Z|\tilde{X}}(t, z)$ satisfying $\pi_{Z|\tilde{X}}(t, z) = \mathbb{P}\left(Z(t) = z \middle| \tilde{X}(s),\ 0 \le s \le t\right)$ almost surely for every $t$. Moreover, since the propensities are uniformly integrable due to Eq. [3], the conditional expectation $\pi_{Z|\tilde{X}}(t, \lambda_j) \triangleq \sum_{z' \in \mathbb{Z}_{\ge 0}^{n_Z}} \lambda_j\left(\tilde{X}(t), z'\right) \pi_{Z|\tilde{X}}(t, z')$ is also cadlag for each $j = 1, \ldots, r$ ([5], Remark 2.27). In the following, we only consider this cadlag conditional probability $\pi_{Z|\tilde{X}}(t, \cdot)$ for the follower system. Also, when discussing other conditional probability, we always refer to such a cadlag one.

Before introducing the equation characterizing the conditional probability $\pi_{Z|\tilde{X}}(t, \cdot)$, we lists a few necessary notations. Following ([6]), we term $\mathcal{O} \triangleq \{j | \zeta_j^{\tilde{X}} \ne \mathbb{0}_{n_{\tilde{X}}}\}$ as the leader-level reactions, where $\mathbb{0}_{n_{\tilde{X}}}$ is the $n_{\tilde{X}}$-dimensional zero vector. Here, we use the notation $\mathcal{O}$ to indicate that these reactions are 'observable' given the trajectory of $\tilde{X}(\cdot)$. Similarly, we term $\mathcal{U} \triangleq \{j | j \notin \mathcal{O}\}$ as the follower-level reactions, where the notation $\mathcal{U}$ indicates that these reactions are 'unobservable' given the trajectory of $\tilde{X}(t)$. Note that for these leader-level reactions, some $\zeta_j^{\tilde{X}}$ might have the same value. To consider this issue, we term

- $\{\xi_1, \ldots, \xi_{r_1}\}$ as the set of non-zero and distinct $\zeta_j^{\tilde{X}}$,
- $\mathcal{O}_{\xi_k} \triangleq \{j | \zeta_j^{\tilde{X}} = \xi_k\}$ $(k = 1, \ldots, r_1)$ as the set in which non-zero $\zeta_j^{\tilde{X}}$ are identical to $\xi_k$,
- $\tilde{R}_{\xi_k}(t) \triangleq \sum_{j \in \mathcal{O}_{\xi_k}} R_j\left(\int_0^t \lambda_j\left(\tilde{X}(s), Z(s)\right) \mathrm{d}s\right)$ as the total firing number of the reactions in $\mathcal{O}_{\xi_k}$ up to time $t$,
- $\lambda^{\mathcal{O}_{\xi_k}}\left(\tilde{X}(s), Z(s)\right) \triangleq \sum_{j \in \mathcal{O}_{\xi_k}} \lambda_j\left(\tilde{X}(s), Z(s)\right)$ as the rate of the process $\tilde{R}_{\xi_k}(t)$,
- $\lambda^{\mathcal{O}}\left(\tilde{X}(s), Z(s)\right) \triangleq \sum_{j \in \mathcal{O}} \lambda_j\left(\tilde{X}(s), Z(s)\right)$.

Note that processes $\tilde{R}_{\xi_1}(t), \ldots, \tilde{R}_{\xi_{r_1}}(t)$ contain the same information as the leader system $\tilde{X}(t)$, meaning that we can construct the former given the trajectory of the latter and vice versa. Then, under the condition Eq. [3], the conditional probability

* $\tilde{\mathcal{X}}_t$ is the filtration generated by the process $X(t)$.

$\pi_{Z|\tilde{X}}(t,z)$ (for any $z \in \mathbb{Z}_{\geq 0}^{n_Z}$), is almost surely characterized by [heuristically derived in (6, 7) and rigorously verified in (8)]

$$
\begin{aligned}
\pi_{Z|\tilde{X}}(t,z) = \pi_{Z|\tilde{X}}(0,z) &+ \int_0^t \sum_{j\in\mathcal{U}} \lambda_j\left(\tilde{X}(s), z-\zeta_j^Z\right)\pi_{Z|\tilde{X}}\left(s, z-\zeta_j^Z\right) - \sum_{j\in\mathcal{U}} \lambda_j\left(\tilde{X}(s), z\right)\pi_{Z|\tilde{X}}(s,z)\mathrm{d}s \\
&- \int_0^t \pi_{Z|\tilde{X}}(s,z)\left(\lambda^{\mathcal{O}}\left(\tilde{X}(s),z\right) - \sum_{z'\in\mathbb{Z}_{\geq 0}^{n_Z}} \lambda^{\mathcal{O}}\left(\tilde{X}(s),z'\right)\pi_{Z|\tilde{X}}(s,z')\right)\mathrm{d}s \\
&+ \sum_{k=1}^{r_1}\int_0^t \left(\frac{\sum_{j\in\mathcal{O}_{\xi_k}} \lambda_j\left(\tilde{X}(s^-), z-\zeta_j^Z\right)\pi_{Z|\tilde{X}}\left(s^-, z-\zeta_j^Z\right)}{\sum_{z'\in\mathbb{Z}_{\geq 0}^{n_Z}} \lambda^{\mathcal{O}_{\xi_k}}\left(\tilde{X}(s^-),z'\right)\pi_{Z|\tilde{X}}(s^-,z')} - \pi_{Z|\tilde{X}}(s^-,z)\right)\mathrm{d}\tilde{R}_{\xi_k}(s),
\end{aligned}
\qquad [14]
$$

and

$$
\int_0^t \sum_{z\in\mathbb{Z}_{\geq 0}^{n_Z}} \sum_{j=1}^{r} \lambda_j(\tilde{X}(s),z)\pi_{Z|\tilde{X}}(s,z)\mathrm{d}s < \infty \text{ almost surely}, \qquad \forall t \geq 0. \qquad [15]
$$

Here, we call Eq. [14] the filtered CME for $Z(t)$.

Now, we show that the follower subsystems $Z_1(t),\ldots,Z_l(t)$ are conditionally independent given the trajectory of the leader system, and the filtered CME Eq. [14] can be divided into several-lower dimensional, independent equations. We first introduce some notations. For each vector $\zeta_j^Z$ ($j = 1,\ldots,r$), we decompose it into $\zeta_j^{Z_1},\ldots,\zeta_j^{Z_l}$, where $\zeta_j^{Z_i}$ corresponds to the state change of $Z_i(t)$ after the firing of the j-th reaction. Also, we denote the dimension of $Z_i(t)$ ($i = 1,\ldots,l$) by $n_{Z_i}$. For each follower subsystem $Z_i(t)$, we term $\mathcal{U}_i \triangleq \left\{j\in\mathcal{U} | \zeta_j^{Z_i} \neq \mathbb{0}_{n_{Z_i}} \text{ or } \lambda_j(\tilde{x}, z_1,\ldots,z_l) \text{ depends on } z_i\right\}$ as the follower-level reactions involving $Z_i(t)$, and we term

$$
\mathcal{O}_i \triangleq \bigcup_{\substack{k:\ \lambda^{\mathcal{O}_{\xi_k}}(\tilde{x},z_1,\ldots,z_l) \text{ depends on } z_i, \\ \text{or } \zeta_j^{Z_i} \neq \mathbb{0}_{n_{Z_i}} \text{for some } j\in\mathcal{O}_{\xi_k}}} \mathcal{O}_{\xi_k}
$$

as the leader-level reactions that involves $Z_i(t)$. According to the conditions C1 and C2, we can easily conclude that

Conclusion 1 The sets $\mathcal{U}_1,\ldots,\mathcal{U}_l$ are disjoint, and $\mathcal{U} = \bigcup_{i=1}^{l}\mathcal{U}$. (By C1 and the definition of $\mathcal{U}_i$)

Conclusion 2 For any $i \in \{1,\ldots,l\}$ and $k \in \{1,\ldots,r_1\}$, there is either $\mathcal{O}_{\xi_k} \subset \mathcal{O}_i$ or $\mathcal{O}_{\xi_k} \cap \mathcal{O}_i = \emptyset$. (By definition)

Conclusion 3 The sets $\mathcal{O}_1,\ldots,\mathcal{O}_l$ are disjoint. (By Conclusion 2 and C2)

Conclusion 4 If $j \in \mathcal{O}/\mathcal{O}_i$, then $\zeta_j^{Z_i} = \mathbb{0}_{n_{Z_i}}$. (By definition).

Then, the conditional independence of the follower subsystems are guaranteed by the following theorem.

**Theorem 1.** *Assume that condition Eq. [3] holds, the follower subsystems $Z_1(t),\ldots,Z_l(t)$ satisfy C1 and C2, and $Z_1(0),\ldots,Z_l(0)$ are conditionally independent given $\tilde{X}(0)$. Then, for any $t > 0$, the follower subsystems $Z_1(t),\ldots,Z_l(t)$ are conditionally independent given the trajectory of $\tilde{X}(\cdot)$ up to time $t$. Their marginal conditional probability defined by $\pi_{Z_i|\tilde{X}}(t,z_i) \triangleq \mathbb{P}\left(Z_i(t) = z_i \middle| \tilde{X}(s),\ 0 \leq s \leq t\right)$ is almost surely characterized by*

$$
\begin{aligned}
\pi_{Z_i|\tilde{X}}(t,z_i) = \pi_{Z_i|\tilde{X}}(0,z_i) &+ \int_0^t \sum_{j\in\mathcal{U}_i} \lambda_j\left(\tilde{X}(s), z_i-\zeta_j^{Z_i}\right)\pi_{Z_i|\tilde{X}}\left(s, z_i-\zeta_j^{Z_i}\right) - \sum_{j\in\mathcal{U}_i} \lambda_j\left(\tilde{X}(s), z_i\right)\pi_{Z_i|\tilde{X}}(s,z_i)ds \\
&- \int_0^t \pi_{Z_i|\tilde{X}}(s,z_i)\left(\lambda^{\mathcal{O}_i}\left(\tilde{X}(s),z_i\right) - \sum_{z_i'\in\mathbb{Z}_{\geq 0}^{n_{Z_i}}} \lambda^{\mathcal{O}_i}\left(\tilde{X}(s),z_i'\right)\pi_{Z_i|\tilde{X}}(s,z_i')\right)ds \\
&+ \sum_{k:\ \mathcal{O}_{\xi_k}\subset\mathcal{O}_i}\int_0^t \left(\frac{\sum_{j\in\mathcal{O}_{\xi_k}} \lambda_j\left(\tilde{X}(s^-), z_i-\zeta_j^{Z_i}\right)\pi_{Z_i|\tilde{X}}\left(s^-, z_i-\zeta_j^{Z_i}\right)}{\sum_{z_i'\in\mathbb{Z}_{\geq 0}^{n_{Z_i}}} \lambda^{\mathcal{O}_{\xi_k}}\left(\tilde{X}(s^-),z_i'\right)\pi_{Z_i|\tilde{X}}(s^-,z_i')} - \pi_{Z_i|\tilde{X}}(s^-,z_i)\right)d\tilde{R}_{\xi_k}(s),
\end{aligned}
\qquad [16]
$$

*and*

$$
\int_0^t \sum_{z_i\in\mathbb{Z}_{\geq 0}^{n_{Z_i}}} \sum_{j\in\mathcal{U}_i\cup\mathcal{O}_i} \lambda_j\left(\tilde{X}(s),z_i\right)\pi_{Z_i|\tilde{X}}(s,z_i)ds < \infty \textit{ almost surely}, \qquad \forall t \geq 0.
$$

*where* $\lambda_j\left(\tilde{X}(t), z_i\right)$ *(for* $j \in \mathcal{U}_i \cup \mathcal{O}_i$*) is the abbreviation of the propensity* $\lambda_j\left(\tilde{X}(t), z_1, \dots, z_l\right)$ *(which does not depend on variables other than* $\tilde{X}(t)$ *and* $z_i$ *due to the definition of* $\mathcal{O}_i$ *and Conclusion 1),* $\lambda^{\mathcal{O}_i}\left(\tilde{X}(s), z'\right) \triangleq \sum_{j\in\mathcal{O}_i} \lambda_j\left(\tilde{X}(s), Z_i(s)\right)$*, and* $n_{Z_i}$ *is the dimension of* $Z_i(t)$*.*

*Proof.* To prove the result, we only need to calculate the conditional probability

$$\pi_{Z_i|\tilde{X}Z_{/i}}(t, z_i) \triangleq \mathbb{P}\left(Z_i(t) = z_i \middle| \tilde{X}(s), Z_1(s), \dots, Z_{i-1}(s), Z_{i+1}(s), \dots, Z_l(s)\ 0 \le s \le t\right)$$

(for $i = 1, \dots, l$) and show it is irrelevant to the trajectory of other follower subsystems, i.e., $Z_1(\cdot), \dots, Z_{i-1}(\cdot), Z_{i+1}(\cdot), \dots, Z_l(\cdot)$. Here, we present the proof for the case $i = 1$; the rest results can be shown in the same way.

To compute $\pi_{Z_i|\tilde{X}Z_{/1}}(t, z_i)$, we can view $\mathcal{U}_1$ as the unobservable reactions given the trajectory of $\tilde{X}(\cdot), Z_2(\cdot), \dots, Z_l(\cdot)$ and the reactions in $\mathcal{O} \cup_{i=2}^{l} \mathcal{U}_i$ as observable reactions given these trajectories. We term

- $\tilde{\mathcal{O}} = \mathcal{O} \cup_{i=2}^{l} \mathcal{U}_i$ as the set of 'observable' reactions in this case,
- $\lambda^{\tilde{\mathcal{O}}/\mathcal{O}_1}(\tilde{x}, z_1, \dots, z_l) \triangleq \sum_{j\in\tilde{\mathcal{O}}/\mathcal{O}_1} \lambda_j(\tilde{x}, z_1, \dots, z_l)$ as the total propensity for the reaction in $\tilde{\mathcal{O}}/\mathcal{O}_1$,
- $\{\eta_1, \dots, \eta_{r_2}\}$ as the set of non-zero and distinct $\left(\zeta_j^{\tilde{X}}, \zeta_j^{Z_2}, \dots, \zeta_j^{l}\right)$ for $j \in \mathcal{O} \cup_{i=2}^{l} \mathcal{U}_i$,
- $\tilde{\mathcal{O}}_{\eta_k} \triangleq \left\{ j \in \mathcal{O} \cup_{i=2}^{l} \mathcal{U}_i \middle| \left(\zeta_j^{\tilde{X}}, \zeta_j^{Z_2}, \dots, \zeta_j^{l}\right) = \eta_k \right\}$ for $k = 1, \dots, r_2$,
- $\tilde{R}_{\eta_k}(t) \triangleq \sum_{j\in\tilde{\mathcal{O}}_{\eta_k}} R_j\left(\int_0^t \lambda_j\left(\tilde{X}(s), Z_1(s), \dots, Z_l(s)\right) \mathrm{d}s\right)$ as the total firing number of the reactions in $\tilde{\mathcal{O}}_{\eta_k}$ up to time $t$,
- $\lambda^{\mathcal{O}_{\eta_k}}\left(\tilde{X}(s), Z_1(s), \dots, Z_l(s)\right) \triangleq \sum_{j\in\mathcal{O}_{\xi_k}} \lambda_j\left(\tilde{X}(s), Z_1(s), \dots, Z_l(s)\right)$ as the rate of the process $\tilde{R}_{\eta_k}(t)$,

Consequently, by Eq. [14], we conclude that $\pi_{Z_1|\tilde{X}Z_{/1}}(t, z_1)$ is characterized by

$$\begin{aligned}
&\pi_{Z_1|\tilde{X}Z_{/1}}(t, z_1) \\
&= \pi_{Z_1|\tilde{X}Z_{/1}}(0, z_1) \\
&\quad + \int_0^t \sum_{j\in\mathcal{U}_1} \lambda_j\left(\tilde{X}(s), z_1 - \zeta_j^{Z_1}, Z_2(s), \dots, Z_l(s)\right) \pi_{Z_1|\tilde{X}Z_{/1}}\left(s, z_1 - \zeta_j^{Z_1}\right) \mathrm{d}s \\
&\quad - \int_0^t \sum_{j\in\mathcal{U}_1} \lambda_j\left(\tilde{X}(s), z_1, Z_2(s), \dots, Z_l(s)\right) \pi_{Z_1|\tilde{X}Z_{/1}}(s, z_1) \mathrm{d}s \\
&\quad - \int_0^t \pi_{Z_1|\tilde{X}Z_{/1}}(s, z_1) \left( \lambda^{\mathcal{O}_1}\left(\tilde{X}(s), z_1, Z_2(s), \dots, Z_l(s)\right) - \sum_{z_1' \in \mathbb{Z}_{\ge 0}^{n_{Z_1}}} \lambda^{\mathcal{O}_1}\left(\tilde{X}(s), z_1', Z_2(s), \dots, Z_l(s)\right) \pi_{Z_1|\tilde{X}Z_{/1}}(s, z_1') \right) \mathrm{d}s \\
&\quad - \int_0^t \pi_{Z_1|\tilde{X}Z_{/1}}(s, z_1) \left( \lambda^{\tilde{\mathcal{O}}/\mathcal{O}_1}\left(\tilde{X}(s), z_1, Z_2(s), \dots, Z_l(s)\right) - \sum_{z_1' \in \mathbb{Z}_{\ge 0}^{n_{Z_1}}} \lambda^{\tilde{\mathcal{O}}/\mathcal{O}_1}\left(\tilde{X}(s), z_1', Z_2(s), \dots, Z_l(s)\right) \pi_{Z_1|\tilde{X}Z_{/1}}(s, z_1') \right) \mathrm{d}s \\
&\quad + \sum_{k:\ \tilde{\mathcal{O}}_{\eta_k} \cap \mathcal{O}_1 \neq \emptyset} \int_0^t \left( \frac{\sum_{j\in\tilde{\mathcal{O}}_{\eta_k}} \lambda_j\left(\tilde{X}(s^-), z_1 - \zeta_j^{Z_1}, Z_2(s), \dots, Z_l(s)\right) \pi_{Z_1|\tilde{X}Z_{/1}}\left(s^-, z_1 - \zeta_j^{Z_1}\right)}{\sum_{z_1' \in \mathbb{Z}_{\ge 0}^{n_{Z_1}}} \lambda^{\tilde{\mathcal{O}}_{\eta_k}}\left(\tilde{X}(s^-), z_1', Z_2(s), \dots, Z_l(s)\right) \pi_{Z_1|\tilde{X}Z_{/1}}(s^-, z_1')} - \pi_{Z_1|\tilde{X}Z_{/1}}(s^-, z_1) \right) \mathrm{d}\tilde{R}_{\eta_k}(s) \\
&\quad + \sum_{k:\ \tilde{\mathcal{O}}_{\eta_k} \cap \mathcal{O}_1 = \emptyset} \int_0^t \left( \frac{\sum_{j\in\tilde{\mathcal{O}}_{\eta_k}} \lambda_j\left(\tilde{X}(s^-), z_1 - \zeta_j^{Z_1}, Z_2(s), \dots, Z_l(s)\right) \pi_{Z_1|\tilde{X}Z_{/1}}\left(s^-, z_1 - \zeta_j^{Z_1}\right)}{\sum_{z_1' \in \mathbb{Z}_{\ge 0}^{n_{Z_1}}} \lambda^{\tilde{\mathcal{O}}_{\eta_k}}\left(\tilde{X}(s^-), z_1', Z_2(s), \dots, Z_l(s)\right) \pi_{Z_1|\tilde{X}Z_{/1}}(s^-, z_1')} - \pi_{Z_1|\tilde{X}Z_{/1}}(s^-, z_1) \right) \mathrm{d}\tilde{R}_{\eta_k}(s),
\end{aligned}$$

which holds almost surely. By the independence condition Eq. [4], we can write $\pi_{Z_1|\tilde{X}Z/1}(0, z_1) = \pi_{Z_1|\tilde{X}}(0, z_1)$. Moreover, By Conclusion 1 and Conclusion 3, the propensities in 3rd to 5th line only depend on the first two arguments. With the same reason, the propensities $\lambda_j(\tilde{x}, z_1, \dots, z_l)$ (for every $j \in \tilde{\mathcal{O}}/\mathcal{O}_1$) depend on $z_1$, and, therefore, the 6th line is zero. Moreover, by Conclusion 2, we can conclude that $\tilde{\mathcal{O}}_{\eta_k} \cap \mathcal{O}_1 \neq \emptyset$ holds only if $\tilde{\mathcal{O}}_{\eta_k} \subset \mathcal{O}_1$; consequently, by Conclusion 4 and the definition of $\mathcal{O}_1$, we can rewrite the 7th line by

$$\sum_{k:\ \tilde{\mathcal{O}}_{\xi_k} \subset \mathcal{O}_1} \int_0^t \left( \frac{\sum_{j\in\tilde{\mathcal{O}}_{\xi_k}} \lambda_j\left(\tilde{X}(s^-), z_1 - \zeta_j^{Z_1}\right) \pi_{Z_1|\tilde{X}Z_{/1}}\left(s^-, z_1 - \zeta_j^{Z_1}\right)}{\sum_{z_1' \in \mathbb{Z}_{\ge 0}^{n_{Z_1}}} \lambda^{\tilde{\mathcal{O}}_{\xi_k}}\left(\tilde{X}(s^-), z_1'\right) \pi_{Z_1|\tilde{X}Z_{/1}}(s^-, z_1')} - \pi_{Z_1|\tilde{X}Z_{/1}}(s^-, z_1) \right) \mathrm{d}\tilde{R}_{\xi_k}(s).$$

By Conclusion 1, Conclusion 3, and Conclusion 4, the propensities in the 8th line does not depend on the second argument, and the vectors $\zeta_j^{Z_1}$ in this line are always zero. Consequently, the 8th line is zero. Finally, by combining all these results, we can rewrite the dynamics of $\pi_{Z_1|\tilde{X}Z/1}(t, z_1)$ (which satisfies Eq. [16]) and find that it is almost surely irrelevant to the trajectory of $Z_2(\cdot), \dots, Z_l(\cdot)$. Moreover, by Eq. [3], $\pi_{Z_1|\tilde{X}Z/1}(t, z_1)$ also satisfies

$$\int_0^t \sum_{z_i \in \mathbb{Z}_{\geq 0}^{n_{Z_i}}} \sum_{j=1}^{r} \lambda_j \left(\tilde{X}(s), Z_1(s), \dots, Z_{i-1}(s), z_i, Z_{i+1}(s), \dots, Z_l(s)\right) \pi_{Z_1|\tilde{X}Z/1}(t, z_1) \mathrm{d}s < \infty \text{ almost surely}, \qquad \forall t \geq 0$$

or equivalently (by Eq. [3])

$$\int_0^t \sum_{z_i \in \mathbb{Z}_{\geq 0}^{n_{Z_i}}} \sum_{j \in \mathcal{U}_i \cup \mathcal{O}_i} \lambda_j \left(\tilde{X}(s), z_i\right) \pi_{Z_i|\tilde{X}}(s, z_i) \mathrm{d}s < \infty \text{ almost surely}, \qquad \forall t \geq 0.$$

Note that the solution of this equation and this inequality is unique, so we can further conclude that $\pi_{Z_1|\tilde{X}Z/1}(t, z_1)$ is also almost surely irrelevant to the trajectory of $Z_2(\cdot), \dots, Z_l(\cdot)$. This implies that the conditional probability $\pi_{Z_1|\tilde{X}}(t, z_1) \triangleq \mathbb{E}\left[\pi_{Z_1|\tilde{X}Z/1}(t, z_1) \middle| \tilde{X}(s),\ 0 \leq s \leq t\right]$ equals to $\pi_{Z_1|\tilde{X}Z/1}(t, z_1)$ almost surely and, therefore, satisfies Eq. [16] and the inequality in this theorem with $i = 1$. $\square$

We call Eq. [16] the filtered CME for the follower subsystem $Z_i(t)$. Also, we can rewrite Eq. [16] into a differential form

$$\begin{aligned} \mathrm{d}\pi_{Z_i|\tilde{X}}(t, z_i) =& f_1\left(\tilde{X}(t), z_i, \pi_{Z_i|\tilde{X}}(t, \cdot)\right) \mathrm{d}t + f_2\left(\tilde{X}(t), z_i, \pi_{Z_i|\tilde{X}}(t, \cdot)\right) \mathrm{d}t \\ &+ \sum_{j=1}^{r} \mathbb{1}\left(\tilde{X}(t) - \tilde{X}(t^-) = \zeta_j^{\tilde{X}}\right) g_j\left(\tilde{X}(t^-), z_i, \pi_{Z_i|\tilde{X}}(t^-, \cdot)\right) \end{aligned} \qquad [17]$$

where

$$f_1\left(\tilde{X}(t), z_i, \pi_{Z_i|\tilde{X}}(t, \cdot)\right) = \sum_{j \in \mathcal{U}_i} \lambda_j\left(\tilde{X}(s), z_i - \zeta_j^{Z_i}\right) \pi_{Z_i|\tilde{X}}\left(s, z_i - \zeta_j^{Z_i}\right) - \sum_{j \in \mathcal{U}_i} \lambda_j\left(\tilde{X}(s), z_i\right) \pi_{Z_i|\tilde{X}}(s, z_i),$$

$$f_2\left(\tilde{X}(t), z_i, \pi_{Z_i|\tilde{X}}(t, \cdot)\right) = -\pi_{Z_i|\tilde{X}}(s, z_i) \left(\lambda^{\mathcal{O}_i}\left(\tilde{X}(s), z_i\right) - \sum_{z_i' \in \mathbb{Z}_{\geq 0}^{n_{Z_i}}} \lambda^{\mathcal{O}_i}\left(\tilde{X}(s), z_i'\right) \pi_{Z_i|\tilde{X}}(s, z_i')\right),$$

and

$$g_j\left(\tilde{X}(t^-), z_i, \pi_{Z_i|\tilde{X}}(t^-, \cdot)\right) = \frac{1}{\left|\mathcal{O}_{\zeta_j^{\tilde{X}}}\right|} \mathbb{1}(j \in \mathcal{O}_i) \sum_{k:\ \xi_k = \zeta_j^{\tilde{X}}} \left(\frac{\sum_{j' \in \mathcal{O}_{\xi_k}} \lambda_{j'}\left(\tilde{X}(s^-), z_i - \zeta_{j'}^{Z_i}\right) \pi_{Z_i|\tilde{X}}\left(s^-, z_i - \zeta_{j'}^{Z_i}\right)}{\sum_{z_i' \in \mathbb{Z}_{\geq 0}^{n_{Z_i}}} \lambda^{\mathcal{O}_{\xi_k}}\left(\tilde{X}(s^-), z_i'\right) \pi_{Z_i|\tilde{X}}(s^-, z_i')} - \pi_{Z_i|\tilde{X}}(s^-, z_i)\right)$$

with $\left|\mathcal{O}_{\zeta_j^{\tilde{X}}}\right|$ the size of $\mathcal{O}_{\zeta_j^{\tilde{X}}}$. In Eq. [17], $f_1$ represents the prediction of the conditional distribution based on the dynamical model, and the remaining terms correspond to the corrections to the estimates in accordance with the dynamics of the observable species.

**B. Derivation of the Rao-Blackwellized CME solver.** We note that for any state $x = (\tilde{x}, z) = (\tilde{x}, z_1, \dots, z_l)$, the probability $p(t, x)$ can be written by

$$p(t, x) = \mathbb{E}\left[\mathbb{1}\left(\tilde{X}(t) = \tilde{x}\right) \pi_{Z|\tilde{X}}(z)\right] = \mathbb{E}\left[\mathbb{1}\left(\tilde{X}(t) = \tilde{x}\right) \prod_{i=1}^{l} \pi_{Z_i|\tilde{X}}(z_i)\right]$$

where the first equality follows from the law of total expectation, and the second follows from Theorem 1. Consequently, we can use the following algorithm to solve the CME master equation.

- Generate $N$ simulations of the system Eq. [2] and denote their leader-system parts by $\tilde{\mathrm{x}}_1(t), \dots, \tilde{\mathrm{x}}_N(t)$.
- For each $\tilde{\mathrm{x}}_j(t)$ and each $Z_i(t)$, we calculate the conditional probability $q_j^i(t, z_i) \triangleq \mathbb{P}\left(Z_i(t) = z_i \middle| \tilde{X}(s) = \tilde{\mathrm{x}}_j(s),\ 0 \leq s \leq t\right)$ using a filtering approach (e.g., the filtered FSP (8))
- Approximate the exact probability by the quantity $\hat{p}_{\mathrm{RB}}(t, x) = \frac{1}{N} \sum_{j=1}^{N} \left[1\left(\tilde{\mathrm{x}}_j(t) = \tilde{x}_j\right) \prod_{i=1}^{l} q_j^i(t, z)\right]$.

We name this algorithm the Rao-Blackwellized CME solver (RB-CME solver).

In this paper's examples, we particular choose the filtered FSP to (8) to solve the filtered CMEs. Specifically, this method approaches the unnormalized version of the filtered CME, which is linear between two neighboring jump time points. This method straightforwardly solves this unnormalized version on a large but finite state space and obtains the filter by normalization. We refer readers to the literature (8) for more details.

**C. Error analysis of the RB-CME solver.** Now, we analyze the error of the RB-CME solver and show that it is more accurate than the Monte-Carlo method given the same sample size $N$. Here, we assume that the error caused by the filtering approach applied to the filtered CME is negligible.

We focus on the $L_1$ error of the RB-CME solver, defined by $\|\hat{p}_{\mathrm{RB}}(t,\cdot)-p(t,\cdot))\|_1 \triangleq \sum_{x\in\mathbb{Z}^n_{\geq 0}} |\hat{p}_{\mathrm{RB}}(t,x)-p(t,x)|$. For every $j\in\{1,\dots,N\}$, $t>0$, and $z=(z_1,\dots,z_l)$, we term $q_j(t,z)=\prod_{i=1}^{l} q_j^i(t,z_i)$ as the conditional probability of the whole follower system given the trajectory $\tilde{\mathrm{x}}_j(\cdot)$ from 0 to $t$. Notice that $\{(\tilde{\mathrm{x}}_j(t),\mathrm{q}_j(t,z))\}_{j=1,\dots,N}$ are independently sampled from the distribution of $\left(\tilde{X}(t),\pi_{Z|\tilde{X}}(t,z)\right)$; therefore, by the law of large numbers, we can conclude

$$\sqrt{N}\left(\hat{p}_{\mathrm{RB}}(t,x)-p(t,x)\right)\xrightarrow{d}\mathcal{N}\left(0,\mathrm{Var}\left(\mathbb{1}\left(\tilde{X}(t)=\tilde{x}\right)\pi_{Z|\tilde{X}}(t,z)\right)\right) \qquad \forall x\in\mathbb{Z}^n_{\geq 0}.$$

We can rewrite the variance by $\mathrm{Var}\left(\mathbb{1}\left(\tilde{X}(t)=\tilde{x}\right)\pi_{Z|\tilde{X}}(t,z)\right)=\mathbb{E}\left[\pi^2_{Z|\tilde{X}}(t,z)\middle|\tilde{X}(t)=\tilde{x}\right]p^{\tilde{X}}(t,\tilde{x})-p^2(t,x)$ where $x=(\tilde{x},z)$, and $p^{\tilde{X}}(t,\tilde{x})=\mathbb{P}\left(\tilde{X}(t)=\tilde{x}\right)$ is the marginal probability of the leader system. Consequently, the error of the RB-CME solver can be written by

$$\lim_{N\to\infty}\sqrt{N}\mathbb{E}\left[\|\hat{p}_{\mathrm{RB}}(t,\cdot)-p(t,\cdot))\|_1\right]=\sqrt{\frac{2}{\pi}}\sum_{x\in\mathbb{Z}^n_{\geq 0}}\sqrt{\mathbb{E}\left[\pi^2_{Z|\tilde{X}}(t,z)\middle|\tilde{X}(t)=\tilde{x}\right]p^{\tilde{X}}(t,\tilde{x})-p^2(t,x)}. \qquad [18]$$

and

$$\lim_{N\to\infty}\sqrt{N}\mathbb{E}\left[\|\hat{p}_{\mathrm{RB}}(t,\cdot)-p(t,\cdot))\|_1\right]=\sqrt{\frac{2}{\pi}}\left[\left(\sum_{x\in\mathbb{Z}^n_{\geq 0}}\sqrt{\mathbb{E}\left[\pi^2_{Z|\tilde{X}}(t,z)\middle|\tilde{X}(t)=\tilde{x}\right]p^{\tilde{X}}(t,\tilde{x})}\right)\pm 1\right]. \qquad [19]$$

Notice that $\pi_{Z|\tilde{X}}(t,z)\in[0,1]$, so we can further conclude

$$\begin{aligned}\lim_{N\to\infty}\sqrt{N}\mathbb{E}\left[\|\hat{p}_{\mathrm{RB}}(t,\cdot)-p(t,\cdot))\|_1\right]&\leq\sqrt{\frac{2}{\pi}}\sum_{x\in\mathbb{Z}^n_{\geq 0}}\sqrt{\mathbb{E}\left[\pi_{Z|\tilde{X}}(t,z)\middle|\tilde{X}(t)=\tilde{x}\right]p^{\tilde{X}}(t,\tilde{x})-p^2(t,x)}\\&\overset{Eq.\ [8]}{=}\lim_{N\to\infty}\sqrt{N}\mathbb{E}\left[\|\hat{p}_{\mathrm{MC}}(t,\cdot)-p(t,\cdot))\|_1\right]\end{aligned}$$

which suggests that the RB-CME solver is no worse than the Monte-Carlo method given a large sample size $N$. In one extreme case where $\pi_{Z|\tilde{X}}(t,z)$ only depends on the final state of $\tilde{X}(\cdot)$ and is irrelevant to its historical information (i.e., $\mathrm{Var}\left[\pi_{Z|\tilde{X}}(t,z)\middle|\tilde{X}(t)=\tilde{x}\right]=0$), we can get

$$\begin{aligned}\lim_{N\to\infty}\sqrt{N}\mathbb{E}\left[\|\hat{p}_{\mathrm{RB}}(t,\cdot)-p(t,\cdot))\|_1\right]&=\sqrt{\frac{2}{\pi}}\sum_{x\in\mathbb{Z}^n_{\geq 0}}\sqrt{\left(\mathbb{E}\left[\pi_{Z|\tilde{X}}(t,z)\middle|\tilde{X}(t)=\tilde{x}\right]\right)^2p^{\tilde{X}}(t,\tilde{x})-p^2(t,x)}\\&=\sqrt{\frac{2}{\pi}}\sum_{\tilde{x}\in\mathbb{Z}^{n_2}_{\geq 0}}\sqrt{p^{\tilde{X}}(t,\tilde{x})\left(1-p^{\tilde{X}}(t,\tilde{x})\right)}.\end{aligned}$$

where the last line is the error of the Monte-Carlo method in estimating the leader system (adapted from Eq. [8]). This formula means that in this extreme case, the RB-CME solver is equivalent to the model reduction method that eliminates all the follower-level species using stochastic averaging; moreover, the RB-CME solver tends to be far more accurate than the Monte-Carlo method in this case, as the leader system usually has a much lower dimension than the whole system by our system decomposition algorithm (see the next subsection). This gives a clue that the RB-CME solvers can be far more accurate than the Monte-Carlo method in general high-dimensional problems.

**D. Automated algorithm for the leader-follower decomposition.** Notice that the decomposition introduced in Section S3.A consists of two parts, where the first one decomposes the system into a leader system and a follower system, and the second part further divides the follower system into several subsystems. Here, we establish the algorithm according to this two-level decomposition strategy.

We first establish the second-level decomposition, i.e., decomposing the whole follower system into several follower subsystems. Notice that this decomposition needs to satisfy C1 and C2, and the maximum size of the follower subsystems should be as small as possible so that the filtering method applied to the filtered CME (see the algorithm of the RB-CME solver) is more efficient. Following these ideas, we propose Algorithm 2 for this decomposition. Specifically, the algorithm first treats each follower-level species as an individual group and then merges them according to the topological conditions C1 and C2. Finally, each group left corresponds to a follower subsystem. One can easily check that this bottom-up algorithm provides the most sparse follower-subsystem decomposition, meaning that any other second-level decomposition satisfying C1 and C2 can only

**Algorithm 2** Second-level decomposition for the RB-CME solver

Input the set of leader-level species and the set of follower-level species. ▷ Input
Classify each follower-level species as an individual group. ▷ Initialization
**for** $j = 1, \dots, r$ **do**
  Merge the groups that involves in the $j$-th reaction. (Denote the merged group by $\texttt{G}_j$) ▷ For C1
**end for**
**for** $k = 1, \dots, r_1$ **do** ▷ For C2
  Merge the groups that have species influencing $\lambda^{\mathcal{O}_{\xi_k}}(x)$. (Denote the merged group by $\tilde{G}_k$)
  **for** $j = 1, \dots, r$ **do**
    **if** $j \in \mathcal{O}_{\xi_k}$ **then**
      Merge $\tilde{G}_k$ with the groups that have species influenced by the $j - th$ reaction.
      $\tilde{G}_k \leftarrow$ the group merged in the previous step.
    **end if**
  **end for**
**end for**
Each group is a follower subsystem. ▷ Output

lead to subsystems that are the union of those provided by Algorithm 2. Moreover, this algorithm has a polynomial complexity with respect to the system size (the species number $n$ and the number of reactions $r$). In general, Algorithm 2 can do the second-level decomposition efficiently and effectively.

We now consider the first-level decomposition, which classifies species into the leader and the follower systems. Since every species can be either a leader-level species or a follower-level one, we have $2^n$ choices in total. Among them, we hope to choose the one that leads to the most accurate RB-CME solver while keeping the solver's time cost below some threshold. As mentioned in the main text, we can interpret this optimal decomposition as the one that maximizes the size of the whole follower system while keeping the size of each individual follower subsystem below a given threshold. Following this idea, we propose Algorithm 3 for the first-level decomposition. Specifically, we first insert a threshold for the maximum size of follower subsystems and provide a truncated state space that contains most probability. After that, we exhaustively search for the optimal decomposition among all the candidates, and finally we use the obtained optimal decomposition to construct the RB-CME solver. Though Algorithm 3 uses the brute-force method, its computational time is acceptable: for a linear network with nine species (Section 2.A in the main text), the algorithm consumes only 3 seconds.

**Algorithm 3** Leader-follower decomposition for the RB-CME solver

Input the threshold $\texttt{T}$ for the maximum size of follower subsystems.
Input a truncated state space $\{0, \dots, \texttt{TS}_1 - 1\} \times \cdots \times \{0, \dots, \texttt{TS}_n - 1\}$ that contains most probability $p(t, x)$. ▷ Input
$\texttt{Largest\_size} \leftarrow 0$. ▷ Initialization
Figure out the $2^n$ choices of the first-level decomposition and give each an index.
**for** $j = 1, \dots, 2^n$ **do** ▷ Search for the optimum
  Use Algorithm 2 to obtain follower subsystems for the $j$-th first-level decomposition candidate.
  $l \leftarrow$ the number of follower subsystems.
  Evaluate the size of each follower subsystem: $\texttt{SS}_i = \prod_{i' \text{: species } S_{i'} \text{ belongs to the } Z_i} \texttt{TS}_{i'}$.
  **if** $\texttt{SS}_i \leq \texttt{T}$ (for all $i \in \{1, \dots, l\}$) and $\prod_{i=1}^{l} \texttt{SS}_i > \texttt{Largest\_size}$ **then** ▷ Check optimality
    Replace the optimal decomposition with the current one.
    $\texttt{Largest\_size} \leftarrow \prod_{i=1}^{l} \texttt{SS}_i$.
  **end if**
**end for**
Output the optimal decomposition. ▷ Output

### S4. Rao-Blackwellized particle filter (RB-PF): derivation and error analysis.

**A. Modified leader-follower decomposition for the filtering problem.** Recall that the filtering problem for intracellular reaction systems can be solved by recursively computing Eq. [6] and Eq. [7], which correspond to prediction and correction, respectively. Following this idea, we propose the Rao-Blackwellized particle filter (RB-PF), which uses the RB-CME solver for the prediction step Eq. [6] and standard techniques for the adjustment step Eq. [7]. Note that the RB-CME solver is established based on the conditional independence among follower subsystems. So, for the RB-PF, this conditional independence needs to be preserved in both the prediction and correction steps. This actually means that the follower subsystems are conditionally independent given the trajectories of both the leader system and the observation process. To this end, we require the following condition for the system decomposition in addition to C1 and C2.

C3 Each observation channel cannot be affected by two follower subsystems. In other words, each entry of $h(\cdot)$ can depend on one follower subsystem at most (besides the leader system).

The theoretical verification is as follows. Let $h_j(\tilde{x}, z_1, \dots, z_1)$ be the $j$-th element of the function $h(x)$. We first term $\mathcal{M}_k \triangleq \{j | h_j(\tilde{x}, z_1, \dots, z_l) \text{ depends on } z_k\}$ as the set of observation channels that measures follower subsystem $Z_k$. By C3, the sets $\mathcal{M}_1, \dots, \mathcal{M}_l$ are disjoint, and the likelihood function can be decomposed by

$$L\left(y|x\right) = L_0(y, \tilde{x}) \prod_{k=1}^{l} L_k(y, \tilde{x}, z_k); \qquad [20]$$

here, $L_0(y, \tilde{x})$ is the density function of the probability $\prod_{j \notin \cup_{k=1}^{l} \mathcal{M}_k} \mathbb{P}\left(Y_j(t_i) \in \cdot \middle| \tilde{X}(t_i) = \tilde{x}\right)$ which can be interpreted as the part of the likelihood irrelevant to the follower subsystems, and $L_k(y, \tilde{x}, z_k)$ is the density function of the probability $\prod_{j \in \mathcal{M}_k} \mathbb{P}\left(Y(t_i) \in \cdot \middle| \tilde{X}(t_i) = \tilde{x}, Z_k(t_i) = z_k\right)$ which can be viewed as the part of the likelihood function relevant to the follower subsystem $Z_k$. With these notations, we can prove the following theorem which guarantees the conditional independence result.

**Theorem 2.** *Under conditions Eq. [3] and Eq. [4], the follower subsystems $Z_1(t), \dots, Z_l(t)$ satisfying C1, C2, and C3 are conditionally independent given the trajectories of $\tilde{X}(\cdot)$ and $Y(\cdot)$ up to time $t$. Moreover, for every $i \in \{0, 1, 2, \dots\}$ and every $t \in (t_i, t_{i+1}]$, the conditional probability $\pi_{Z_k|\tilde{X}Y}(t, z_k) \triangleq \mathbb{P}\left(Z_k(t) = z_k \middle| \tilde{X}(s),\ 0 \le s \le t, \text{ and } Y(t_j),\ 0 \le j \le i\right)$ is almost surely characterized by Eq. [16] with the initial condition $\mathbb{P}\left(Z_k(t_i) = z_k \middle| \tilde{X}(s)\ 0 \le s \le t_i, \text{ and } Y(t_j),\ 0 \le j \le i\right)$ and*

$$\int_0^t \sum_{z_k \in \mathbb{Z}_{\ge 0}^{n_{Z_k}}} \sum_{j \in \mathcal{U}_i \cup \mathcal{O}_i} \lambda_j\left(\tilde{X}(s), z_k\right) \pi_{Z_k|\tilde{X}Y}(t, z_k) ds < \infty \text{ almost surely}, \qquad \forall t \ge 0.$$

*Proof.* We prove the rest result by mathematical induction. For $t = 0$, the result holds immediately from Theorem 1. Then, we assume that the theorem holds for $t \in [0, t_i]$ and check whether it holds for $t \in [t_i, t_{i+1}]$.

Now, we show that for all $t \in (t_i, t_{i+1}]$, the process $\pi_{Z_k|\tilde{X}Y}(t, z_k) \triangleq \mathbb{P}\left(Z_k(t) = z_k \middle| \tilde{X}(s),\ 0 \le s \le t, \text{ and } Y(t_j),\ 0 \le j \le i\right)$ is almost surely characterized by Eq. [16] with the initial condition $\mathbb{P}\left(Z_k(t_i) = z_k \middle| \tilde{X}(s)\ 0 \le s \le t_i, \text{ and } Y(t_j),\ 0 \le j \le i\right)$. Note that $\pi_{Z_k|\tilde{X}Y}(t, z_k) = \mathbb{E}\left[\pi_{Z_k|\tilde{X}}(t, z_k) \middle| \tilde{X}(s),\ 0 \le s \le t, \text{ and } Y(t_j),\ 0 \le j \le i\right]$. By applying this relation to Theorem 1, we can show that for all $t \in [t_i, t_{i+1}]$, the conditional probability $\pi_{Z_k|\tilde{X}Y}(t, z_k)$ almost surely satisfies Eq. [16] with the initial condition $\mathbb{P}\left(Z_k(t_i) = z_k \middle| \tilde{X}(s)\ 0 \le s \le t_i, \text{ and } Y(t_j),\ 0 \le j \le i\right)$. Also, note that the solution is unique given the inequality in the theorem (see Theorem 1).

By this characterization equation and the conditional independence of $Z_1(t_i), \dots, Z_l(t_i)$ given $\{\tilde{X}(s),\ 0 \le s \le t_i\}$ and $\{Y(t_j),\ 0 \le j \le i\}$, we can say that the conditional probability $\pi_{Z_k|\tilde{X}Y}(t, z_k)$ (for every $k = 1, \dots, l$ and $t \in [t_i, t_{i+1}]$) is almost surely irrelevant to the trajectories of the follower subsystems other than $Z_k$. Therefore, for $t \in [t_i, t_{i+1}]$, the follower subsystems $Z_1(t), \dots, Z_l(t)$ are conditionally independent given $\{\tilde{X}(s),\ 0 \le s \le t\}$ and $\{Y(t_j),\ 0 \le j \le i\}$.

Finally, we prove that the follower subsystem are still conditionally independent after the collection of the observation at time $t_{i+1}$. By Bayes' rule and Eq. [20], we have

$$\begin{aligned}
&\mathbb{P}\left(Z_1(t_{i+1}) = z_1, \dots, Z_l(t_{i+1}) = z_l \middle| \tilde{X}(s),\ 0 \le s \le t_{i+1}, \text{ and } Y(t_j),\ 0 \le j \le i+1\right) \\
&\propto \prod_{k=1}^{l} L_k\left(Y(t_{i+1}), \tilde{X}(t_{i+1}), z_k\right) \mathbb{P}\left(Z_k(t_{i+1}) = z_k \middle| \tilde{X}(s),\ 0 \le s \le t_{i+1}, \text{ and } Y(t_j),\ 0 \le j \le i\right) \\
&= \prod_{k=1}^{l} \mathbb{P}\left(Z_k(t_{i+1}) = z_k \middle| \tilde{X}(s),\ 0 \le s \le t_{i+1}, \text{ and } Y(t_j),\ 0 \le j \le i+1\right)
\end{aligned}$$

where the last line follows from the definition of the conditional marginal distribution. This suggests that $Z_1(t_{i+1}), \dots, Z_l(t_{i+1})$ are conditionally independent given $\{\tilde{X}(s),\ 0 \le s \le t_{i+1}\}$ and $\{Y(t_j),\ j = 1, \dots, i+1\}$.

Combining all the above results, we show the theorem for $t \in [t_i, t_{i+1}]$. Moreover, by math induction, we prove the result. □

Now, we modify the leader-follower decomposition algorithm according to C3. Note that this additional requirement only affects the second-level decomposition, which divides the follower system into subsystems, and does not affect the first-level decomposition, which classifies species into the leader and follower systems according to an optimization rule. Following this idea, we propose a modified second-level decomposition algorithm in Algorithm 4. Compared with Algorithm 2 (used for the CME solver), this new algorithm additionally puts fluorescent reporters of the same color into the same subsystem (see the 16th line). Then, we provide the whole leader-follower decomposition algorithm for the RB-PF in Algorithm 5, which uses Algorithm 4 rather than Algorithm 3 for the second-level decomposition (see the 6th line).

**Algorithm 4** Second-level decomposition for the RB-PF

```
1:  Input the set of leader-level species and the set of follower-level species.            ▷ Input
2:  Classify each follower-level species as an individual group.                            ▷ Initialization
3:  for j = 1, ..., r do
4:      Merge the groups that are involved in the j-th reaction. (Denote the merged group by G_j)   ▷ For C1
5:  end for
6:  for k = 1, ..., r_1 do                                                                    ▷ For C2
7:      Merge the groups that have species influencing λ^{O_{ξ_k}}(x). (Denote the merged group by G̃_k)
8:      for j = 1, ..., r do
9:          if j ∈ O_{ξ_k} then
10:             Merge G̃_k with the groups that have species influenced by the j − th reaction.
11:             G̃_k ← the group merged in the previous step.
12:         end if
13:     end for
14: end for
15: Merge the groups that have the same color fluorescent reporters.                          ▷ For C3
16: Each group is a follower subsystem.                                                       ▷ Output
```

**Algorithm 5** Leader-follower decomposition for the RB-PF

```
1:  Input the threshold T for the maximum size of follower subsystems.
2:  Input a truncated state space {0, ..., TS_1 − 1} × ··· × {0, ..., TS_n − 1} that contains most probability.   ▷ Input
3:  Largest_size ← 0.                                                                         ▷ Initialization
4:  Figure out the 2^n choices of the first-level decomposition and give each an index.
5:  for j = 1, ..., 2^n do                                                                    ▷ Search for the optimum
6:      Use Algorithm 4 to obtain follower subsystems for the j-th first-level decomposition candidate.
7:      l ← the number of follower subsystems.
8:      Evaluate the size of each follower subsystem: SS_i = ∏_{i': species S_{i'} belongs to the Z_i} TS_{i'}.
9:      if SS_i ≤ T (for all i ∈ {1, ..., l}) and ∏_{i=1}^{l} SS_i > Largest_size then         ▷ Check optimality
10:         Replace the optimal decomposition with the current one.
11:         Largest_size ← ∏_{i=1}^{l} SS_i.
12:     end if
13: end for
14: Output the optimal decomposition.                                                         ▷ Output
```

**B. Algorithm of the RB-PF.** By Theorem 2, we can rewrite the prediction probability $\rho_{t_{i+1}}(\cdot)$ and the filter $\pi_{t_{i+1}}(\cdot)$ by

$$\rho_{t_{i+1}}(\tilde{x}, z_1, \ldots, z_l) = \mathbb{E}\left[ \mathbb{1}\left(\tilde{X}(t_{i+1}) = \tilde{x}\right) \prod_{k=1}^{l} \pi_{Z_k|\tilde{X}Y}(t_{i+1}, z_k) \middle| Y(t_j),\ 0 \le j \le i \right] \qquad [21]$$

$$\pi_{t_{i+1}}(\tilde{x}, z_1, \ldots, z_l) = \mathbb{E}\left[ \mathbb{1}\left(\tilde{X}(t_{i+1}) = \tilde{x}\right) \prod_{k=1}^{l} \bar{\pi}_{Z_k|\tilde{X}Y}(t_{i+1}, z_k) \middle| Y(t_j),\ 0 \le j \le i+1 \right] \qquad [22]$$

where $\pi_{Z_k|\tilde{X}Y}(t_{i+1}, z_k) \triangleq \mathbb{P}\left(Z_k(t_{i+1}) = z_k \middle| \tilde{X}(s),\ 0 \le s \le t_{i+1},\ \text{and } Y(t_j),\ 0 \le j \le i\right)$ and

$$\begin{aligned}\bar{\pi}_{Z_k|\tilde{X}Y}(t_{i+1}, z_k) &\triangleq \mathbb{P}\left(Z_k(t_{i+1}) = z_k \middle| \tilde{X}(s),\ 0 \le s \le t_{i+1},\ \text{and } Y(t_j),\ 0 \le j \le i+1\right) \\ &\propto L_k\left(Y(t_{i+1}), \tilde{X}(t_{i+1}), z_k\right) \pi_{Z_k|\tilde{X}Y}(t_{i+1}, z_k). \qquad \text{(by Bayes' rule and Eq. [20])}\end{aligned}$$

Theorem 2 also states that the conditional probability $\pi_{Z_k|\tilde{X}Y}(t, z_k)$ (for $t \in (t_i, t_{i+1}]$) satisfies the filtered CME Eq. [16] with the initial probability $\bar{\pi}_{Z_k|\tilde{X}Y}(t_i, z_k)$. Consequently, we can solve the filtering problem by applying the RB-CME solver to the prediction probability $\rho_{t_{i+1}}(\cdot)$ in Eq. [21] (or equivalently Eq. [6]).

Following this idea, we provide the detailed algorithm of the RB-PF in Algorithm 6. Specifically, this algorithm first decomposes the system into a leader system and several follower subsystems using Algorithm 5. Then, in each iteration, the algorithm computes the prediction probability $\rho_{t_{i+1}}(\cdot)$ by the RB-CME solver (see line 6–10). The prediction probability is approximated by $\hat{\rho}^{\mathrm{RB}}_{t_{i+1}}(x) = \sum_{j=1}^{N} w_j(t)\mathbb{1}\left(\tilde{\mathrm{x}}_j(t_{i+1}) = \tilde{x}\right)\prod_{k=1}^{l} \bar{\mathrm{q}}_j^k(t_{i+1}, z_k)$ where $w_j(t)$ is the weight of the $j$-th particle, $\tilde{\mathrm{x}}_j(t_{i+1})$ is the leader part of the $j-th$ particle (simulated in Line 6–8), and $\mathrm{q}_j^k(t_{i+1}, z_k)$ is the conditional probability of the subsystem $Z_k$ given $\tilde{\mathrm{x}}_j(\cdot)$ and $Y(\cdot)$ up to time $t_{i+1}$. Here, $\hat{\rho}^{\mathrm{RB}}_{t_{i+1}}(x)$ is not explicitly given in the algorithm because it is not necessary. After the prediction step, the RB-PF updates the weights for each $\tilde{\mathrm{x}}_j(t_{i+1})$ according to the adjustment step Eq. [7] and reevaluates the conditional probability of follower subsystems by including new observations (line 11–12). Finally, Algorithm 6 computes the filter according to Eq. [22] (line 13) and resamples the particles to delete non-significant ones (line 14).

**Algorithm 6** Rao-Blackwellized particle filter (RB-PF)

1: Decompose the systems using Algorithm 5. ▷ System decomposition
2: Generate $N$ samples $(\mathrm{x}_1(0), \dots, \mathrm{x}_N(0))$ from the initial probability and give them equal weights $(w_j(0) = \frac{1}{N}, j = 1, \dots, N)$.
3: Denote the leader parts of these particles by $(\tilde{\mathrm{x}}_1(0), \dots, \tilde{\mathrm{x}}_N(0))$. For each $\tilde{\mathrm{x}}_j(0)$, each subsystem $Z_k$, and each $z_k$ in a finite state space, set $\bar{\mathrm{q}}_j^k(0, z_k) \triangleq \mathbb{P}\left(Z_k(0) = z_k \middle| \tilde{X}(0) = \tilde{\mathrm{x}}_k(0)\right)$. ▷ Initialization
4: $i \leftarrow 0$ and $t_0 \leftarrow 0$.
5: **while** $t_i$ is not the final observation time **do**
6: Construct particles: for each $j \in \{1, \dots, N\}$ and each subsystems $Z_k$, sample a $\tilde{z}_k(t_i)$ from the distribution $\bar{q}_j^k(t_i, \cdot)$;
7: then the $j$-th particle becomes $(\tilde{\mathrm{x}}_j(t_i), \tilde{z}_1(t_i), \dots, \tilde{z}_\ell(t_i))$
8: Simulate every particle $\mathrm{x}_j(t)$ from time $t_i$ to $t_{i+1}$ according to Eq. [2] and denote their leader parts by $\tilde{\mathrm{x}}_1(\cdot), \dots, \tilde{\mathrm{x}}_N(\cdot)$.
9: For each $\tilde{\mathrm{x}}_j(\cdot)$ and $Z_k$, use a filtering approach (i.e., the filtered FSP (8)) to solve Eq. [16] from time $t_i$ to $t_{i+1}$ with the initial probability $\bar{\mathrm{q}}_j^k(t_i, z_k)$.
10: Denote the solution at $t_{i+1}$ by $\mathrm{q}_j^k(t_{i+1}, z_k)$ ▷ Prediction (Eq. [6] and Eq. [21])
11: Update weights $w_j(t_{i+1}) \propto w_j(t_i)L_0(Y(t_{i+1}), \tilde{\mathrm{x}}_j(t_{i+1}))\prod_{k=1}^{l}\left(\sum_{z_k} L_k\left(Y(t_{i+1}), \tilde{\mathrm{x}}_j(t_{i+1}), z_k\right)\mathrm{q}_j^k(t_{i+1}, z_k)\right)$.
12: $\bar{\mathrm{q}}_j^k(t_{i+1}, z_k) \propto L_k\left(Y(t_{i+1}), \tilde{\mathrm{x}}_j(t_{i+1}), z_k\right)\mathrm{q}_j^k(t_{i+1}, z_k)$. ▷ Adjustment (Eq. [7] and Eq. [22])
13: Compute the filter $\hat{\pi}^{\mathrm{RB}}_{t_{i+1}}(x) = \sum_{j=1}^{N} w_j(t_{i+1})\mathbb{1}\left(\tilde{\mathrm{x}}_j(t_{i+1}) = \tilde{x}\right)\prod_{k=1}^{l}\bar{\mathrm{q}}_j^k(t_{i+1}, z_k)$.
14: Resample $\left\{w_j(t_{i+1}), \left(\tilde{\mathrm{x}}_j(t_{i+1}), \bar{\mathrm{q}}_j^1(t_{i+1}, \cdot), \dots, \bar{\mathrm{q}}_j^l(t_{i+1}, \cdot)\right)\right\}$ to obtain N equally weighted particles
15: $i \leftarrow i + 1$.
16: **end while**

**C. Error analysis of the RB-PF.** Now, we analyze the error of the RB-PF. First, we investigate the error at the first observation time point $t_1$ and show that its performance (for the filtering problem) is similar to that of the RB-CME solver (for solving the CME). Again, we assume that the error of the filtering approach applied to the filtered CME is negligible. Readers who are not interested in the detailed derivation can directly go to Eq. [24].

For the $j$-th particle, we term $\mathrm{q}_j(t_1, z) \triangleq \prod_{i=1}^{l} \mathrm{q}_j^l(t_1, z_i)$ as the conditional distribution of $Z(t_1)$ given the trajectory of the leader system $\tilde{\mathrm{x}}_j(\cdot)$. Then, we can rewrite the RB-PF filter at $t_1$ by

$$\hat{\pi}^{\mathrm{RB}}_{t_1}(x) = \frac{\frac{1}{N}\sum_{j=1}^{N}\left(\frac{L\left(Y(t_1)\middle|\left(\tilde{\mathrm{x}}_j(t_1), z\right)\right)}{\rho^Y_{t_1}(Y(t_1))}\right)\mathbb{1}\left(\tilde{\mathrm{x}}_j(t_1) = \tilde{x}\right)\mathrm{q}_j(t_1, z)}{\frac{1}{N}\sum_{j=1}^{N}\sum_{z'}\left(\frac{L\left(Y(t_1)\middle|\left(\tilde{\mathrm{x}}_j(t_1), z'\right)\right)}{\rho^Y_{t_1}(Y(t_1))}\right)\mathrm{q}_j(t_1, z')} = \frac{\frac{1}{N}\sum_{j=1}^{N}\left(\frac{L\left(Y(t_1)\middle|\left(\tilde{\mathrm{x}}_j(t_1), z\right)\right)}{\rho^Y_{t_1}(Y(t_1))}\right)\mathbb{1}\left(\tilde{\mathrm{x}}_j(t_1) = \tilde{x}\right)\mathrm{q}_j(t_1, z)}{\frac{1}{N}\sum_{j=1}^{N}\left(\frac{q_j^Y(t_1, Y(t_1))}{\rho^Y_{t_1}(Y(t_1))}\right)\mathrm{q}_j(t_1, z)}$$

where $\rho^Y_{t_1}(y) \triangleq \sum_{\tilde{x}} L(y|x)\rho_{t_1}(x)$ is the probability density of $Y(t_1)$, and $q_j^Y(t_1, y) \triangleq \sum_{z'} L\left(y|(\tilde{\mathrm{x}}_j(t_1), z')\right)q_j(t_1, z')$ is the estimate of $Y(t_1)$'s probability density by the $j$-th particle. By applying the delta method to this formula, we have

$$\sqrt{N}\left[\hat{\pi}^{\mathrm{RB}}_{t_1}(\tilde{x}, z) - \pi_{t_1}(\tilde{x}, z)\right] \xrightarrow{d} \mathcal{N}\left(0, \mathrm{Var}\left(\frac{L\left(Y(t_1)\,|(\tilde{\mathrm{x}}_1(t_1), z)\right)}{\rho^Y_{t_1}\left(Y(t_1)\right)}\mathbb{1}\left(\tilde{\mathrm{x}}_1(t_1) = \tilde{x}\right)\mathrm{q}_1(t_1, z) - \frac{q_1^Y\left(t_1, Y(t_1)\right)}{\rho^Y_{t_1}\left(Y(t_1)\right)}\pi_{t_1}(\tilde{x}, z)\middle| Y(t_1)\right)\right).$$

given $Y(t_1)$ and, therefore,

$$\lim_{N\to\infty}\sqrt{N}\mathbb{E}\left[\left|\hat{\pi}^{\mathrm{RB}}_{t_1}(\tilde{x}, z) - \pi_{t_1}(\tilde{x}, z)\right|\middle| Y(t_1)\right] = \sqrt{\frac{2}{\pi}}\sqrt{\mathrm{Var}\left(\frac{L\left(Y(t_1)\,|(\tilde{\mathrm{x}}_1(t_1), z)\right)}{\rho^Y_{t_1}\left(Y(t_1)\right)}\mathbb{1}\left(\tilde{\mathrm{x}}_1(t_1) = \tilde{x}\right)\mathrm{q}_1(t_1, z) - \frac{q_1^Y\left(t_1, Y(t_1)\right)}{\rho^Y_{t_1}\left(Y(t_1)\right)}\pi_{t_1}(\tilde{x}, z)\middle| Y(t_1)\right)}. \quad [23]$$

Since the particle $(\tilde{\mathrm{x}}_1(t_1), \mathrm{q}_1(t_1, z))$ has the same distribution as $\left(\tilde{X}(t_1), \pi_{Z|\tilde{X}}(t_1, z)\right)$ and is independent of $Y(t_1)$, the conditional variance in Eq. [23] satisfies

$$
\begin{aligned}
&\mathrm{Var}\left( \frac{L\left(Y(t_1)\,|(\tilde{\mathrm{x}}_1(t_1), z)\,\right)}{\rho^Y_{t_1}\left(Y\left(t_1\right)\right)} \mathbb{1}\left(\tilde{\mathrm{x}}_1(t_1) = \tilde{x}\right) \mathrm{q}_1(t_1, z) - \frac{q^Y_1\left(t_1, Y(t_1)\right)}{\rho^Y_{t_1}\left(Y\left(t_1\right)\right)} \pi_{t_1}(\tilde{x}, z) \middle| Y(t_1) \right) \\
&\leq \left( \frac{L\left(Y(t_1)\,|(\tilde{x}, z)\,\right)}{\rho^Y_{t_1}\left(Y\left(t_1\right)\right)} \right)^2 \rho^{\tilde{X}}_{t_1}(\tilde{x}) \mathbb{E}\left[ \pi^2_{Z|\tilde{X}}(t_1, z) \middle| \tilde{X}(t_1) = \tilde{x} \right] + \frac{\mathbb{E}\left[ \left(q^Y_1\left(t_1, Y(t_1)\right)\right)^2 \middle| Y(t_1) \right]}{\left(\rho^Y_{t_1}\left(Y\left(t_1\right)\right)\right)^2} \pi^2_{t_1}(\tilde{x}, z) \\
&\leq \left( \frac{L\left(Y(t_1)\,|(\tilde{x}, z)\,\right)}{\rho^Y_{t_1}\left(Y\left(t_1\right)\right)} \right)^2 \rho^{\tilde{X}}_{t_1}(\tilde{x}) \mathbb{E}\left[ \pi^2_{Z|\tilde{X}}(t_1, z) \middle| \tilde{X}(t_1) = \tilde{x} \right] \\
&\quad + \frac{\mathbb{E}\left[ \sum_{z'} L^2\left(y|(\tilde{\mathrm{x}}_j(t_1), z')\right) q_j(t_1, z') \middle| Y(t_1) \right]}{\left(\rho^Y_{t_1}\left(Y\left(t_1\right)\right)\right)^2} \pi^2_{t_1}(\tilde{x}, z) \qquad \text{(Jensen's inequality)} \\
&\leq \left( \frac{L\left(Y(t_1)\,|(\tilde{x}, z)\,\right)}{\rho^Y_{t_1}\left(Y\left(t_1\right)\right)} \right)^2 \rho^{\tilde{X}}_{t_1}(\tilde{x}) \mathbb{E}\left[ \pi^2_{Z|\tilde{X}}(t_1, z) \middle| \tilde{X}(t_1) = \tilde{x} \right] + \left( \frac{\det(\sqrt{2\pi\sigma})^{-1/2}}{\rho^Y_{t_1}\left(Y\left(t_1\right)\right)} \right) \pi^2_{t_1}(\tilde{x}, z) \qquad \left(L\left(y|x\right) \leq \det(\sqrt{2\pi\sigma})^{-1/2}\right)
\end{aligned}
$$

where $\rho^{\tilde{X}}_{t_1}(\tilde{x}) \triangleq \sum_z \rho_{t_1}(\tilde{x}, z)$ is the marginal distribution of $\tilde{X}(t_1)$, and

$$
\begin{aligned}
&\mathrm{Var}\left( \frac{L\left(Y(t_1)\,|(\tilde{\mathrm{x}}_1(t_1), z)\,\right)}{\rho^Y_{t_1}\left(Y\left(t_1\right)\right)} \mathbb{1}\left(\tilde{\mathrm{x}}_1(t_1) = \tilde{x}\right) \mathrm{q}_1(t_1, z) - \frac{q^Y_1\left(t_1, Y(t_1)\right)}{\rho^Y_{t_1}\left(Y\left(t_1\right)\right)} \pi_{t_1}(\tilde{x}, z) \middle| Y(t_1) \right) \\
&\geq \mathbb{E}\left[ \left( \frac{L\left(Y(t_1)\,|(\tilde{\mathrm{x}}_1(t_1), z)\right)}{\rho^Y_{t_1}\left(Y\left(t_1\right)\right)} \right)^2 \mathbb{1}\left(\tilde{\mathrm{x}}_1(t_1) = \tilde{x}\right) \mathrm{q}^2_1(t_1, z) - 2 \times \mathbb{1}\left(\tilde{\mathrm{x}}_1(t_1) = \tilde{x}\right) \frac{L\left(Y(t_1)\,|(\tilde{\mathrm{x}}_1(t_1), z)\,\right) \mathrm{q}_1(t_1, z)}{\rho^Y_{t_1}\left(Y\left(t_1\right)\right)} \frac{q^Y_1\left(t_1, Y(t_1)\right)}{\rho^Y_{t_1}\left(Y\left(t_1\right)\right)} \pi_{t_1}(\tilde{x}, z) \middle| Y(t_1) \right] \\
&\geq \mathbb{E}\left[ \left( \frac{L\left(Y(t_1)\,|(\tilde{\mathrm{x}}_1(t_1), z)\,\right)}{\rho^Y_{t_1}\left(Y\left(t_1\right)\right)} \right)^2 \mathbb{1}\left(\tilde{\mathrm{x}}_1(t_1) = \tilde{x}\right) \mathrm{q}^2_1(t_1, z) - 2 \times \mathbb{1}\left(\tilde{\mathrm{x}}_1(t_1) = \tilde{x}\right) \frac{\pi_{t_1}(\tilde{x}, z) \mathrm{q}_1(t_1, z)}{\rho_{t_1}(\tilde{x}, z)} \frac{\det(\sqrt{2\pi\sigma})^{-1/2}}{\rho^Y_{t_1}\left(Y\left(t_1\right)\right)} \pi_{t_1}(\tilde{x}, z) \middle| Y(t_1) \right] \\
&= \left( \frac{L\left(Y(t_1)\,|(\tilde{x}, z)\,\right)}{\rho^Y_{t_1}\left(Y\left(t_1\right)\right)} \right)^2 \rho^{\tilde{X}}_{t_1}(\tilde{x}) \mathbb{E}\left[ \pi^2_{Z|\tilde{X}}(t_1, z) \middle| \tilde{X}(t_1) = \tilde{x} \right] - 2 \left( \frac{\det(\sqrt{2\pi\sigma})^{-1/2}}{\rho^Y_{t_1}\left(Y\left(t_1\right)\right)} \right) \pi^2_{t_1}(\tilde{x}, z).
\end{aligned}
$$

By applying these results together with the relation $\sqrt{a+b} \leq \sqrt{a} + \sqrt{b}$ to the formula Eq. [23], we can estimate the $L_1$ error of the RB-PF filter by

$$
\begin{aligned}
&\lim_{N\to\infty} \sqrt{N} \mathbb{E}\left[ \left\| \hat{\pi}^{\mathrm{RB}}_{t_1}(\cdot) - \pi_{t_1}(\cdot) \right\|_1 \right] \\
&\triangleq \lim_{N\to\infty} \sum_{(\tilde{x}, z) \in \mathbb{Z}^n_{\geq 0}} \sqrt{N}\ \mathbb{E}\left[ \left| \hat{\pi}^{\mathrm{RB}}_{t_1}(\tilde{x}, z) - \pi_{t_1}(\tilde{x}, z) \right| \right] \\
&= \sqrt{\frac{2}{\pi}} \left[ \left( \sum_{x \in \mathbb{Z}^n_{\geq 0}} \sqrt{\rho^{\tilde{X}}_{t_1}(\tilde{x}) \mathbb{E}\left[ \pi^2_{Z|\tilde{X}}(t_1, z) \middle| \tilde{X}(t_1) = \tilde{x} \right]} \right) \pm 2 \det(\sqrt{2\pi\sigma})^{-1/4} \int_{y \in \mathbb{R}^m} \sqrt{\rho^Y_{t_1}(y)}\, \mathrm{d}y \right] \qquad [24]
\end{aligned}
$$

provided that both terms in the last line are convergent. Here, $m$ is the dimension of $Y(t_1)$.

The formula Eq. [24] tells that the error of the RB-PF at time $t_1$ largely depends on the two quantities in the square brackets. The first term in this bracket represents the error of the RB-CME solver in estimating the prediction probability $\rho_{t_1}(\cdot)$ (see Eq. [19]), and the second term shows the dispersion of $\rho^Y_{t_1}(\cdot)$ which tends to grow exponentially with $m$. Note that the observation dimension $m$ is usually very low, and the observation noise is not negligible. So, we can conclude that the term $\sum_{x \in \mathbb{Z}^n_{\geq 0}} \sqrt{\rho^{\tilde{X}}_{t_1}(\tilde{x}) \mathbb{E}\left[ \pi^2_{Z|\tilde{X}}(t_1, z) \middle| \tilde{X}(t_1) = \tilde{x} \right]}$ usually dominates in Eq. [24]; in other words, the performance of the RB-PF (for the filtering problem) is similar to that of the RB-CME solver (for solving CMEs). This result suggests that if one system's CME can be accurately solved by the RB-CME solver, then its filtering problem can also be accurately solved by the RB-PF and vice versa. Also, since the RB-CME solver scales favorably with the system dimension (as discussed in the main text), the RB-PF should also scale favorably with the system dimension.

From Eq. [13] and the discussion after that, we can observe that this performance consistency also applies to the Monte-Carlo method (for solving the CME) and the particle filter (for the filtering problem). Therefore, we conjecture that this consistency is a universal property for CME solvers and their associated filters.

For observation time points other than $t_1$, we conjecture that the same results should also hold. The theoretical analysis can be performed in the same way as we presented above, except that the particles $\left(\tilde{\mathrm{x}}_j(t_i), q^1_j(t_i, \cdot), \dots, q^l_j(t_i, \cdot)\right)$ have a slightly different distribution from that of $\left(\tilde{X}(t_i), \pi_{Z_1|\tilde{X}Y}(t_i, \cdot), \dots, \pi_{Z_l|\tilde{X}Y}(t_i, \cdot)\right)$ due to the adjustment step and resampling. A rigorous analysis for that requires the techniques in (4), which is somehow complicated. However, we conjecture that this discrepancy should not affect the performance of the RB-PF too much, and, therefore, the results obtained for time $t_1$ should also apply to other time points. Our numerical studies also support this point (see Section 2.C in the main text). We leave the theoretical verification for further work.

### S5. Rao-Blackwell method for cell-specific model identification: derivation and algorithms.

**A. Problem statement.** Now, we introduce the mathematics for model identification, whose main difference from before is the incorporation of parameter uncertainty. We still consider a chemical reaction network Eq. [1], which consists of all possible reactions in the considered cell. Similar to Eq. [2], the dynamical equation can be written by

$$X(t) = X(0) + \sum_{j=1}^{r} \zeta_j R_j \left( \int_0^t \lambda_j(\Theta, X(s)) \mathrm{d}s \right) \qquad [25]$$

where $\Theta$ is an $\tilde{r}$-vector of model parameters (e.g., reaction constant and hill coefficients) taking values in a discrete state space $\mathbb{O} \subset \mathbb{R}^{\tilde{r}}_{\geq 0}$, and the remaining terms have the same meaning as the ones in Eq. [2]. Compared with Eq. [2], the propensity function $\lambda_j(\cdot)$ in Eq. [25] has an additional dependence on model parameters, thereby accounting for parameter uncertainty. Similar to Eq. [3], we consider a non-explosivity condition:

$$\sum_{j=1}^{n} \mathbb{E}\left[\lambda_j^2(\Theta, X(t))\right] \text{ is uniformly bounded on any time interval } [0,T], \qquad [26]$$

This condition also implies that the random variables $\{\lambda_j(\Theta, X(t))\}_{t\in[0,T]}$ are uniformly integrable. Moreover, similar to Eq. [4], we assume that all the parameters and the initial conditions for different species are independent, i.e.,

$$\text{all the elements in } \Theta \text{ and } X(0) \text{ are independent} \qquad [27]$$

We still considered the experiments under fluorescent microscope platforms as in the filtering case, where a cell is measured at different time points $\{t_1, \dots, t_{n_f}\}$ with measurements $Y(t_i)$ satisfying Eq. [5]. Model identification aims to calculate the conditional distribution of $\Theta$ given all the measurements, i.e., $\mathbb{P}\left(\Theta = \cdot \;|Y(t_s), 1 \leq s \leq t_{\mathrm{n}_f}\right)$. By Bayes' rule, the solution of this identification problem can be solved by a series of recursive formulas:

$$\rho_{t_{i+1}}(\theta, x) = \sum_{x\in\mathbb{Z}^n_{\geq 0}} \mathbb{P}\left(X(t_{i+1}) = x \middle| \Theta = \theta, X(t_i) = x'\right) \pi_{t_i}(\theta, x') \qquad \text{for } i = 0, 1, \dots, \mathrm{n}_f - 1 \qquad [28]$$

$$\pi_{t_{i+1}}(\theta, x) \propto L\left(Y(t_{i+1})|x\right) \rho_{t_{i+1}}(\theta, x) \qquad \text{for } i = 0, 1, \dots, \mathrm{n}_f - 1 \qquad [29]$$

$$\mathbb{P}\left(\Theta = \theta \;|Y(t_s), 1 \leq s \leq t_{\mathrm{n}_f}\right) = \sum_{x\in\mathbb{Z}^n_{\geq 0}} \pi_{t_{\mathrm{n}_f}}(\theta, x) \qquad [30]$$

where $\pi_{t_i}(\theta, x) \triangleq \mathbb{P}(\Theta = \theta, X(t_i) = x|Y(t_s), 1 \leq s \leq i)$ and $\rho_{t_{i+1}}(\theta, x) \triangleq \mathbb{P}(\Theta = \theta, X(t_{i+1}) = x|Y(t_s), 1 \leq s \leq i)$.

**B. Connection between model identification and stochastic filtering.** Essentially, we can view $(\Theta, X(t))$ as a state of an expanded chemical reaction network, where $\Theta$ represents some additional special chemical species that can take non-integer values and remain constant over time. From this viewpoint, the probability distribution of $(\Theta, X(t))$ (denoted by $p(t, \theta, x) \triangleq \mathbb{P}(\Theta = \theta, X(t) = x)$) follows an augmented CME:

$$\frac{\mathrm{d}p(t,\theta,x)}{\mathrm{d}t} = \sum_{j=1}^{r} \lambda_j(\theta, x - \zeta_j) p(t, \theta, x - \zeta_j) - \sum_{j=1}^{r} \lambda_j(\theta, x) p(t, \theta, x), \qquad \forall \theta \in \mathbb{O}, \text{ and } \forall x \in \mathbb{Z}_{\geq 0}.$$

Moreover, model identification can be viewed as a type of filtering problem that aims to infer the additional hidden states $\Theta$. This point is also reflected in the formulas Eq. [28] and Eq. [29], where Eq. [28] solves an augmented CME with the initial condition $\pi_{t_i}(\cdot)$, and Eq. [29] adjusts the prediction $\rho_{t_{i+1}}(\cdot)$ according to the new observation $Y(t_{i+1})$.

**C. Rao-Blackwell method for model identification.** Due to the similarities between model identification and stochastic filtering, the RB-PF introduced earlier can be adapted to address model identification challenges. The RB-PF consists of two fundamental components: 1) leader-follower decomposition and 2) the Rao-Blackwell algorithm for calculating the conditional distribution. In the subsequent discussion, we will explore how to modify these components for the use in model identification context.

***C.1. Modification of the leader-follower decomposition.*** The RB-PF (introduced earlier) utilizes a hybrid approach to infer different components of the system, where the leader part is inferred using particle filtering, and the follower part is inferred with the assistance of a filtering approach. In this paper, we particularly choose the filtered FSP (8) as the filtering approach. Classical particle filtering is known to be inefficient for inferring static hidden variables (e.g., model parameters) due to sample degeneracy, where the efficient sample size for the inference of these static variables drops dramatically over time (9). Though several modification methods have been developed to address this issue (such as the resample-move method (10, 11), regularized particle filtering (9, 12, 13) and nested particle filtering (14, 15)), they all introduce extra noise into the inference algorithm, necessitating fine-tuning to strike a balance between sample degeneracy and the added noise, which can be extremely time-consuming (13). Consequently, we choose to classify all the parameters as part of the follower component of the system to circumvent sample degeneracy and avoid additional noise.

C4 All the model parameters $\Theta$ are classified as follower components of the system.

Following C4, we can decompose the whole system state $(\Theta, X(t))$ into a leader system $\tilde{X}(t)$, and several follower subsystems $(\Theta_1, Z_1(t)), \ldots, (\Theta_\ell, Z_\ell(t))$, where the dimension of a particular $\Theta_i$ or $Z_i(t)$ $(i = 1, \ldots, \ell)$ can be zero, but the dimensions of $\Theta_i$ and $Z_i(t)$ cannot be zero simultaneously. We term $\mathbb{O}_i$ as the feasible region of $\Theta_i$. To indicate the contribution of model parameters to the propensity function, we redefine $\mathcal{U}_i \triangleq \left\{ j \in \mathcal{U} | \zeta_j^{Z_i} \neq \mathbb{0}_{n_{Z_i}} \text{ or } \lambda_j(\theta_1, \ldots, \theta_\ell, \tilde{x}, z_1, \ldots, z_\ell) \text{ depends on } (\theta_i, z_i) \right\}$, which represents follower-level reactions involving $(\Theta_i, Z_i(t))$, and

$$\mathcal{O}_i \triangleq \bigcup_{\substack{k:\ \ \lambda^{\mathcal{O}_{\xi_k}}(\theta_1, \ldots, \theta_\ell, \tilde{x}, z_1, \ldots, z_\ell) \text{ has dependence on } \theta_i \text{ or } z_i, \\ \text{or } \zeta_j^{Z_i} \neq \mathbb{0}_{n_{Z_i}} \text{ for some } j \in \mathcal{O}_{\xi_k}}} \mathcal{O}_{\xi_k}$$

which represents the leader-level reactions that involves $(\Theta_i, Z_i(t))$. All other notations introduced previously are kept unchanged.

Recall that the model parameters can be viewed as static chemical species (see S5.C); therefore, we can get a result similar to Theorem 2 based on Conditions C1 — C3. Specifically, if the follower subsystems satisfy these conditions, then they are conditionally independent given the trajectory of the leader system and observations, and the dynamics of their conditional probability distributions have expressions similar to Eq. [16]. More details are given in the following theorem.

**Theorem 3** (Adapted from Theorem 2)**.** *Under conditions Eq. [26] and Eq. [27], the follower subsystems* $(\Theta_1, Z_1(t)), \ldots, (\Theta_\ell, Z_\ell(t))$ *satisfying C1, C2, and C3 are conditionally independent given the trajectory of* $\tilde{X}(\cdot)$ *and* $Y(\cdot)$ *up to time* $t$*. Moreover, for every* $k \in \{0, 1, 2, n_f - 1\}$ *and every* $t \in (t_{\tilde{k}}, t_{\tilde{k}+1}]$*, the conditional probability*

$$\pi_{\Theta_i Z_i | \tilde{X} Y}(t, \theta_i, z_i) \triangleq \mathbb{P}\left(\Theta_i = \theta_i, Z_i(t) = z_i \middle| \tilde{X}(s),\ 0 \leq s \leq t,\ \textit{and } Y(t_j),\ 0 \leq j \leq \tilde{k}\right)$$

*is almost surely characterized by*

$$\begin{aligned}
&\pi_{\Theta_i Z_i | \tilde{X} Y}(t, \theta_i, z_i) \\
&= \mathbb{P}\left(\Theta_i = \theta_i, Z_i(t_i) = z_i \middle| \tilde{X}(s),\ 0 \leq s \leq t_i,\ \textit{and } Y(t_j),\ 0 \leq j \leq i\right) \\
&\quad + \int_{t_{\tilde{k}}}^{t} \sum_{j \in \mathcal{U}_i} \lambda_j\left(\theta_i, \tilde{X}(s), z_i - \zeta_j^{Z_i}\right) \pi_{\Theta_i Z_i | \tilde{X} Y}\left(s, \theta_i, z_i - \zeta_j^{Z_i}\right) - \sum_{j \in \mathcal{U}_i} \lambda_j\left(\theta_i, \tilde{X}(s), z_i\right) \pi_{\Theta_i Z_i | \tilde{X} Y}(s, \theta_i, z_i)\, ds \\
&\quad - \int_{t_{\tilde{k}}}^{t} \pi_{\Theta_i Z_i | \tilde{X} Y}(s, \theta_i, z_i) \left( \lambda^{\mathcal{O}_i}\left(\theta_i, \tilde{X}(s), z_i\right) - \sum_{(\theta_i', z_i') \in \mathbb{O}_i \times \mathbb{Z}_{\geq 0}^{n_{Z_i}}} \lambda^{\mathcal{O}_i}\left(\theta_i', \tilde{X}(s), z_i'\right) \pi_{\Theta_i Z_i | \tilde{X} Y}\left(s, \theta_i', z_i'\right) \right) ds \\
&\quad + \sum_{k:\ \mathcal{O}_{\xi_k} \subset \mathcal{O}_i} \int_0^t \left( \frac{\sum_{j \in \mathcal{O}_{\xi_k}} \lambda_j\left(\theta_i, \tilde{X}(s^-), z_i - \zeta_j^{Z_i}\right) \pi_{\Theta_i Z_i | \tilde{X} Y}\left(s^-, \theta_i, z_i - \zeta_j^{Z_i}\right)}{\sum_{(\theta_i', z_i') \in \mathbb{O}_i \times \mathbb{Z}_{\geq 0}^{n_{Z_i}}} \lambda^{\mathcal{O}_{\xi_k}}\left(\theta_i', \tilde{X}(s^-), z_i'\right) \pi_{\Theta_i Z_i | \tilde{X} Y}\left(s^-, \theta_i', z_i'\right)} - \pi_{\Theta_i Z_i | \tilde{X} Y}\left(s^-, \theta_i, z_i\right) \right) d\tilde{R}_{\xi_k}(s),
\end{aligned} \tag{31}$$

*and*

$$\int_0^t \sum_{(\theta_i', z_i') \in \mathbb{O}_i \times \mathbb{Z}_{\geq 0}^{n_{Z_i}}} \sum_{j \in \mathcal{U}_i \cup \mathcal{O}_i} \lambda_j\left(\theta_i, \tilde{X}(s), z_i, \right) \pi_{\Theta_i Z_i | \tilde{X} Y}(s, \theta_i, z_i) ds < \infty \ \textit{a.s. for all } t \in [t_{\tilde{k}}, t_{\tilde{k}+1}].$$

*Proof.* Note that the model parameters can be viewed as static chemical species (see S5.C). Therefore, this theorem is a straightforward consequence of Theorem 2. □

This theorem together with Theorem 2 demonstrate that the leader-follower decompositions for stochastic filtering and model identification bear a considerable resemblance to each other. The leader-follower decomposition for model identification requires further consideration only of the involvement of parameters in each reaction and the fulfillment of condition C4, which stipulates that the parameters should be classified as follower components. From this perspective, we modify the two level decomposition algorithms (Algorithm 4 and Algorithm 5) to obtain new ones for model identification (see Algorithm 7 and Algorithm 8). Specifically, the second-level decomposition additionally considers the involvement of model parameters (see Lines 2, 4, and 7 in Algorithm 7); meanwhile, the first-level decomposition classify only species as leader components (see Line 5 in Algorithm 8) and include the size of parameter space when calculating the size of follower subsystems (see Line 9 in Algorithm 8).

**Algorithm 7** Second-level decomposition in the Rao-Blackwell method for model identification

```
Input the set of leader-level species and the set of follower-level species.                      ▷ Input
Classify each follower-level species and model parameter as an individual group.          ▷ Initialization
for j = 1, ..., r do
    Merge the groups that are involved in the j-th reaction. (Denote the merged group by G_j)   ▷ For C1
end for
for k = 1, ..., r_1 do                                                                            ▷ For C2
    Merge the groups that have species and parameters influencing λ^{O_{ξ_k}}(x). (Denote the merged group by G̃_k)
    for j = 1, ..., r do
        if j ∈ O_{ξ_k} then
            Merge G̃_k with the groups that have species influenced by the j − th reaction.
            G̃_k ← the group merged in the previous step.
        end if
    end for
end for
Merge the groups that have the same color fluorescent reporters.                                  ▷ For C3
Each group is a follower subsystem.                                                               ▷ Output
```

**Algorithm 8** Leader-follower decomposition for model identification

```
Input the threshold T for the maximum size of follower subsystems.
Input a truncated state space for species {0, ..., TS_1 − 1} × ··· × {0, ..., TS_n − 1} that contains most probability.
Input the number of admissible values for each model parameter, represented by S^θ_1, ..., S^θ_r̃        ▷ Input
Largest_size ← 0.                                                                             ▷ Initialization
Figure out the 2^n choices of the first-level decomposition and give each an index.
for j = 1, ..., 2^n do                                                                 ▷ Search for the optimum
    Use Algorithm 7 to obtain follower subsystems for the j-th first-level decomposition candidate.
    l ← the number of follower subsystems.
    Evaluate the size of each follower subsystem: SS_i = (∏_{i': i'−th paremter belongs to Θ_i} S^θ_{i'}) (∏_{i': species S_{i'} belongs to the Z_i} TS_{i'}).
    if SS_i ≤ T (for all i ∈ {1, ..., l}) and ∏_{i=1}^{l} SS_i > Largest_size then          ▷ Check optimality
        Replace the optimal decomposition with the current one.
        Largest_size ← ∏_{i=1}^{l} SS_i.
    end if
end for
Output the optimal decomposition.                                                                 ▷ Output
```

***C.2. Rao-Blackwell algorithm for model identification.*** Now, we present the detailed algorithm applying Rao-Blackwell method to model identification. Similar to the filtering problem, the key of the algorithm is to represent the prediction and correction steps (Eq. [28] and Eq. [29]) in a Rao-Blackwell format. By Theorem 3, the prediction probability $\rho_{t_{i+1}}(\cdot)$ can be re-written by

$$\rho_{t_{i+1}}(\theta_1,\dots,\theta_\ell,\tilde{x},z_1,\dots,z_\ell) = \mathbb{E}\left[\mathbb{1}\left(\tilde{X}(t_{i+1})=\tilde{x}\right)\prod_{k=1}^{\ell}\pi_{\Theta_k Z_k|\tilde{X}Y}(t_{i+1},\theta_k,z_k)\middle| Y(t_j),\ 0\le j\le i\right] \quad [32]$$

where $\pi_{\Theta_k Z_k|\tilde{X}Y}(t_{i+1},\theta_k,z_k)$ is given in Theorem 3, and the adjusted probability $\pi_{t_{i+1}}(\cdot)$ can be rewritten by

$$\pi_{t_{i+1}}(\theta_1,\dots,\theta_\ell,\tilde{x},z_1,\dots,z_\ell) = \mathbb{E}\left[\mathbb{1}\left(\tilde{X}(t_{i+1})=\tilde{x}\right)\prod_{k=1}^{l}\bar{\pi}_{\Theta_k Z_k|\tilde{X}Y}(t_{i+1},\theta_k,z_k)\middle| Y(t_j),\ 0\le j\le i+1\right] \quad [33]$$

where $\bar{\pi}_{\Theta_k Z_k|\tilde{X}Y}(t_{i+1},\theta_k,z_k) \triangleq \mathbb{P}\left(\Theta_k=\theta_k, Z_k(t_{i+1})=z_k \middle| \tilde{X}(s),\ 0\le s\le t_{i+1},\ \text{and}\ Y(t_j),\ 0\le j\le i+1\right)$ and satisfies

$$\bar{\pi}_{\Theta_k Z_k|\tilde{X}Y}(t_{i+1},\theta_k,z_k) \propto L_k\left(Y(t_{i+1}),\tilde{X}(t_{i+1}),z_k\right)\pi_{\Theta_k Z_k|\tilde{X}Y}(t_{i+1},\theta_k,z_k) \qquad \text{(by Bayes' rule and Eq. [20])}.$$

Also, Theorem 3 states that the term $\pi_{\Theta_k Z_k|\tilde{X}Y}(t,\theta_k,z_k)$ (for $t\in(t_i,t_{i+1})$) satisfies Eq. [31] with the initial condition $\bar{\pi}_{\Theta_k Z_k|\tilde{X}Y}(t_i,\theta_k,z_k)$. With these formulas, we can construct a Rao-Blackwell method for model identification (see Algorithm 9). Essentially, this algorithm for model identification follows the same structure as the RB-PF (refer to Algorithm 6), but it also takes model parameters into account. Again, this is not surprising as the model parameters can be viewed as static chemical species, and the model identification problem with time-course data has a strong resemblance to the stochastic filtering problem (see S5.C).

**Algorithm 9** Rao-Blackwell method for model identification (with time-course data)

Decompose the systems using Algorithm 8. ▷ System decomposition

Sample $N$ species state $\mathrm{x}_1(0), \ldots, \mathrm{x}_N(0)$ from the initial probability and give them equal weights ($w_j(0) = \frac{1}{N}$, $j = 1, \ldots, N$).

Denote the leader parts of these samples by $(\tilde{\mathrm{x}}_1(0), \ldots, \tilde{\mathrm{x}}_N(0))$. For each $\tilde{\mathrm{x}}_j(0)$, each subsystem $(\Theta_k, Z_k)$, and each pair $(\theta_k, z_k)$, set $\bar{\mathrm{q}}_j^k(0, \theta_k, z_k) \triangleq \mathbb{P}\left(\Theta_k = \theta_k, Z_k(0) = z_k \middle| \tilde{X}(0) = \tilde{\mathrm{x}}_k(0)\right)$. ▷ Initialization

$i \leftarrow 0$ and $t_0 \leftarrow 0$.

**while** $t_i$ is not the final observation time **do**

  Construct particles: for each $j \in \{1, \ldots, N\}$ and each subsystems $(\theta_k, z_k)$, sample a pair $\left(\tilde{\theta}_k, \tilde{z}_k(t_i)\right)$ from the distribution $\bar{q}_j^k(t_i, \cdot)$; then the $j$-th particle becomes $(\tilde{\theta}_1, \ldots, \tilde{\theta}_\ell, \tilde{\mathrm{x}}_j(t_i), \tilde{z}_1(t_i), \ldots, \tilde{z}_\ell(t_i))$

  Simulate every particle from time $t_i$ to $t_{i+1}$ according to Eq. [25] and denote the leader parts by $\tilde{\mathrm{x}}_1(\cdot), \ldots, \tilde{\mathrm{x}}_N(\cdot)$.

  For each $\tilde{\mathrm{x}}_j(\cdot)$ and each subsystem $(\Theta_k, Z_k(t))$, use the filtered FSP to solve Eq. [31] in $[t_i, t_{i+1}]$ with the initial condition $\bar{\mathrm{q}}_j^k(t_i, \theta_k, z_k)$. Denote the solution at $t_{i+1}$ by $\mathrm{q}_j^k(t_{i+1}, \theta_k, z_k)$ ▷ Prediction (Eq. [6] and Eq. [32])

  Update weights $w_j(t_{i+1}) \propto w_j(t_i) L_0(Y(t_{i+1}), \tilde{\mathrm{x}}_j(t_{i+1})) \prod_{k=1}^{\ell} \left(\sum_{z_k} L_k\left(Y(t_{i+1}), \tilde{\mathrm{x}}_j(t_{i+1}), z_k\right) \mathrm{q}_j^k(t_{i+1}, \theta_k, z_k)\right)$.

  $\bar{\mathrm{q}}_j^k(t_{i+1}, \theta_k, z_k) \propto L_k\left(Y(t_{i+1}), \tilde{\mathrm{x}}_j(t_{i+1}), z_k\right) \mathrm{q}_j^k(t_{i+1}, \theta_k, z_k)$. ▷ Adjustment (Eq. [7] and Eq. [33])

  Compute the filter $\hat{\pi}_{t_{i+1}}^{\mathrm{RB}}(\theta, x) = \sum_{j=1}^{N} w_j(t_{i+1}) \mathbb{1}\left(\tilde{\mathrm{x}}_j(t_{i+1}) = \tilde{x}\right) \prod_{k=1}^{\ell} \bar{\mathrm{q}}_j^k(t_{i+1}, \theta_k, z_k)$.

  Resample $\left\{w_j(t_{i+1}), \left(\tilde{\mathrm{x}}_j(t_{i+1}), \bar{\mathrm{q}}_j^1(t_{i+1}, \cdot), \ldots, \bar{\mathrm{q}}_j^\ell(t_{i+1}, \cdot)\right)\right\}$ to obtain N equally weighted particles

  $\mathrm{i} = \mathrm{i} + 1$

**end while**

### S6. Modeling of the genetic circuits in cases studies

**A. Modeling of the repressilator.** Here, we introduce the details of the repressilator model presented in the main text. First, we built the model according to the literature (16). Specifically, this model consists of three gene expression systems producing cI, lacI, and tetR, respectively, and the protein products cyclically repress each other's expression. The involved chemical reactions are listed as followers, where $S_1, S_2, \ldots, S_6$ represents the cI mRNA, cI, lacI mRNA, lacI, tetR mRNA, and tetR, respectively.

$$
\begin{array}{llll}
\emptyset \xrightarrow{\frac{\phi_1}{\left(1+(X_6(t))^2\right)}+\phi_2} S_1 & \text{Transcription} & S_1 \xrightarrow{k_2} \emptyset & \text{Degradation} \\
S_1 \xrightarrow{k_3} S_1 + S_2 & \text{Translation} & S_2 \xrightarrow{k_4} \emptyset & \text{Degradation} \\
\emptyset \xrightarrow{\frac{\phi_1}{\left(1+(X_2(t))^2\right)}+\phi_2} S_3 & \text{Transcription} & S_3 \xrightarrow{k_6} \emptyset & \text{Degradation} \\
S_3 \xrightarrow{k_7} S_3 + S_4 & \text{Translation} & S_4 \xrightarrow{k_8} \emptyset & \text{Degradation} \\
\emptyset \xrightarrow{\frac{\phi_1}{\left(1+(X_4(t))^2\right)}+\phi_2} S_5 & \text{Transcription} & S_5 \xrightarrow{k_{10}} \emptyset & \text{Degradation} \\
S_5 \xrightarrow{k_{11}} S_5 + S_6 & \text{Translation} & S_6 \xrightarrow{k_{12}} \emptyset & \text{Degradation}
\end{array}
$$

In these reactions, the first two lines correspond to the gene expression system of cI, the third and fourth lines correspond to the gene expression system of lacI, and the last two lines correspond to the gene expression system of tetR. The values of model parameters and the initial conditions are listed in Table S1.

| Model parameter $(\text{min}^{-1})$ | Initial condition |
|---|---|
| $k_2 = 0.3$ | $X_1(0) = 1$ |
| $k_3 = 2$ | $X_2(0) = 50$ |
| $k_4 = 0.07$ | $X_3(0) = 0$ |
| $k_6 = 0.3$ | $X_4(0) = 0$ |
| $k_7 = 2$ | $X_5(0) = 0$ |
| $k_8 = 0.07$ | $X_6(0) = 0$ |
| $k_{10} = 0.3$ | |
| $k_{11} = 2$ | |
| $k_{12} = 0.07$ | |
| $\phi_1 = 0.5$ | |
| $\phi_2 = 0.5 \times 10^{-4}$ | |

**Table S1. Performance of**

In this example, we solve the associated CME at time 500, before which a few oscillations have taken place. Through a few simulations, we found that most of the probability is contained in the state space where each mRNA has fewer than 20 copies, and each protein has fewer than 200 copies. (The simulations are not shown in the paper.) Note that this state space contains 64 billion states, and storing a probability on this state space requires 476.8 GB (8 bytes for each state). Consequently, the FSP is impractical for this problem. To have an accurate approximation of the exact probability, we simulated $3 \times 10^9$ trajectories of the system and viewed the empirical distribution as the exact probability. The whole procedure took 10 GPUs ($3 \times 10^8$ simulations for each) about 24 hours! Recall that storing the whole probability is impractical, so we only stored the marginal distribution of the mRNAs and the marginal distribution of the proteins. We also applied the RB-CME solver and the Monte-Carlo method to this example, and all the results are shown in the main text. The performance of the RB-CME solver and Monte-Carlo method is evaluated by the sum of the $L_1$ errors in estimating the marginal distributions of the mRNAs and proteins.

**B. Modeling of the genetic toggle switch.** Here, we introduce the details of the genetic toggle switch presented in the main text. This circuit consists of two gene expression systems whose protein products repress each other's expression. Following the literature (17), we model the genetic toggle switch by the following reactions, where $S_1$ is the activated state of the first gene, $S_2$ is the deactivated state of the first gene, $S_3$ is the protein product of the first gene, $S_4$ is the activated state of the second gene, $S_5$ is the deactivated state of the second gene, and $S_6$ is the protein product of the second gene.

$$\begin{array}{llll}
S_1 \xrightarrow{k_1} S_1 + S_3 & \text{(Gene expression)} & S_4 \xrightarrow{k_2} S_4 + S_6 & \text{(Gene expression)} \\
S3 \xrightarrow{k_3} \emptyset & \text{(Degradation )} & S_6 \xrightarrow{k_4} \emptyset & \text{(Degradation)} \\
S_2 \xrightarrow{k_5} S_1 & \text{(Activation} & S_5 \xrightarrow{k_6} S_4 & \text{(Activation)} \\
S_1 \xrightarrow{\phi_2 \exp(\phi_1 X_6(t))} S_2 & \text{(Deactivation)} & S_4 \xrightarrow{\phi_3 \exp(\phi_1 X_3(t))} S_5 & \text{(Deactivation)}
\end{array}$$

In these reactions, the first column corresponds to the first gene expression system, and the second column corresponds to the second gene expression system. The model parameters and initial conditions are presented in Table S2. Moreover, we assume

| Model parameter $\left(\text{min}^{-1}\right)$ | Initial condition |
|---|---|
| $k_1 = 60$ | $X_1(0) = 0$ |
| $k_2 = 60$ | $X_2(0) = 1$ |
| $k_3 = 0.5$ | $X_3(0) = 0$ |
| $k_4 = 0.5$ | $X_4(0) = 1$ |
| $k_5 = 0.3$ | $X_5(0) = 0$ |
| $k_6 = 0.3$ | $X_6(0) = 20$ |
| $\phi_1 = 0.05$ | |
| $\phi_2 = 0.1$ | |
| $\phi_3 = 0.1$ | |

**Table S2. Model parameters and initial conditions of the genetic toggle switch**

that the first protein is fluorescent and measured by a microscope every 20 minutes. Moreover, we assume the observation to satisfy

$$Y(t_i) = X_3(t_i) + \sigma W_i, \qquad \forall t_i \in \{20, 40, \ldots, 200\}, \tag{34}$$

where $\{t_i\}$ are the observation time points, $\sigma$ is the observation noise intensity, and $\{W_i\}$ is a sequence of independent standard Gaussian white noise.

In this example, our goal is to estimate the hidden dynamical state based on these partial observations. Through a few simulations, we found that most of the probability is contained in the state space where each protein has fewer than 200 copies. Then, we applied the FSP with this specific state space to the associated filtering problem and viewed its solution as the ground truth. Also, we applied both the RB-PF and the PF to the problem and depicted their errors by the distances between their solutions and the exact filter. The numerical results are presented in the main text.

### S7. Modeling of Transcription system in Yeast Cells

**A. Reaction network model for this transcription system.** The transcription system consists of four species: $G_0$ (inactive gene), $G_1$ (active gene), $G_2$ (gene in an advanced active state), and mRNA (messenger RNA). We represent their respective molecular counts by $X_{G_0}$, $X_{G_1}$, $X_{G_2}$, and $X_{\text{mRNA}}$. Then, the chemical reactions occurring within this system are modeled as follows.

$$
\begin{array}{lll}
G_0 \xrightarrow{k_1} G_1 & \text{(Initial activation of the gene)} & \text{Propensity: } k_1 X_{G_0}(t) \\
G_1 \xrightarrow{k_2} G_0 & \text{(Deactivation of the gene)} & \text{Propensity: } k_2 X_{G_1}(t) \\
G_1 \xrightarrow{k_3} G_2 & \text{(Full activation of the gene)} & \text{Propensity: } k_3 X_{G_1}(t) \\
G_2 \xrightarrow{k_4} G_1 & \text{(Partial deactivation of the gene)} & \text{Propensity: } k_4 X_{G_2}(t) \\
\emptyset \longrightarrow \text{mRNA} & \text{(Transcription)} & \text{Propensity: } k_{p1} X_{G_1}(t) + (k_{p1} + kp2) X_{G_2}(t) \\
\text{mRNA} \longrightarrow \emptyset & \text{(Degradation of mRNAs)} & \text{Propensity: } X_{\text{mRNA}}
\end{array}
$$

Here, the gene transitions among three gene states $G_0$, $G_1$, and $G_2$, and the mRNA are transcribed when the gene is in either of the two active states, $G_1$ and $G_2$. At the initial time point, the gene is in the state $G_0$, and the mRNA count equals zero.

Moreover, the cell is measured under a microscope at different time points $\{t_1, \dots, t_{n_f}\}$, and the measurements satisfy the relation

$$Y(t_i) = X_{\text{mRNA}}(t_i) + \sigma W_i$$

where $\sigma$ is the noise intensity, and $\{W_i\}_{i=1,\dots,n_f}$ are independent standard Gaussian random variables.

For the ease of notation, let us denote $G_0$, $G_1$, $G_2$, and mRNA as $S_1$, $S_2$, $S_3$, and $S_4$, respectively. Similarly, we denote $X_{G_0}$, $X_{G_1}$, $X_{G_2}$, and $X_{\text{mRNA}}$ as $X_1$, $X_2$, $X_3$, and $X_4$.

**B. Ergodicity of the system.** Now, we show that the system exhibits ergodicity in some invariant state space when reasonable parameters are chosen. Recall that given an invariant space, ergodicity means the occupation time distribution $\mathbb{P}_{oc}(T, x) \triangleq \frac{1}{T}\int_0^T \mathbb{1}\left(X(t) = x\right) \mathrm{d}t$ almost surely converges to the stationary distribution on this space as $T \to \infty$. The following analysis are based on the method in (18).

First, we consider the case where all the parameters are positive. In this case, we can observe that the system state $(X_1(t), X_2(t), X_3(t), X_4(t))$ evolves in an invariant space $\mathcal{IS} \triangleq \left\{x \in \mathbb{Z}^4_{\geq 0} | x_1 + x_2 + x_3 = 1\right\}$, and the stochastic system is irreducible on this state space. Furthermore, by denoting $v = (1, 1, 1, 1)^\top$, we have that

$$
\begin{aligned}
\sum_{j=1}^{6} \lambda_j(\theta, x) < v, \zeta_j > &= k_{p1}x_2 + (k_{p1} + k_{p2})x_3 - x_4 \leq (k_{p1} + k_{p2}) + 1 - < v, x > && \forall x \in \mathcal{IS} \\
\sum_{j=1}^{6} \lambda_j(\theta, x) < v, \zeta_j >^2 &= k_{p1}x_2 + (k_{p1} + k_{p2})x_3 + x_4 \leq (k_{p1} + k_{p2}) + < v, x > && \forall x \in \mathcal{IS}.
\end{aligned}
$$

Therefore, according to (18, Proposition 4), the system has a unique stationary distribution on $\mathcal{IS}$ for any positive parameters, and the system also exhibits ergodicity.

Second, we consider the scenario where the system has only one active gene state, meaning that $k_1$, $k_2$, and $k_{p1}$ are positive, but $k_3 = 0$. In this case, we can also find an invariant measure $\mathcal{IS}_{x_3=0} \triangleq \left\{x \in \mathbb{Z}^4_{\geq 0} | x_1 + x_2 = 1, x_3 = 0\right\}$, and the stochastic system is irreducible on this state space. Moreover, by denoting $v = (1, 1, 1, 1)^\top$, we have that

$$
\begin{aligned}
\sum_{j=1}^{6} \lambda_j(\theta, x) < v, \zeta_j > &= k_{p1}x_2 - x_4 \leq k_{p1} + 1 - < v, x > && \forall x \in \mathcal{IS}_{x_3=0} \\
\sum_{j=1}^{6} \lambda_j(\theta, x) < v, \zeta_j >^2 &= k_{p1}x_2 + x_4 \leq k_{p1} + < v, x > && \forall x \in \mathcal{IS}_{x_3=0}.
\end{aligned}
$$

Consequently, according to (18, Proposition 4), the system is also ergodic when $k_1, k_2, k_{p1} > 0$ but $k_3 = 0$.

### S8. Noise decomposition based on ergodicity

Now, we utilize the stability of the underlying continuous-time Markov chain, i.e. ergodicity, to propose a method for noise decomposition from single-cell time-lapse microscopy data. We consider that a microscopic platform can record the dynamics of a particular chemical species in individual cells. This dynamics is denoted by $\{X_\theta(t)\}_{t\geq 0}$ with $\theta$ the system parameters that can be different from cell-to-cell due to *extrinsic variability.* Also, We assume that for each cell the value of $\theta$ is constant and unchanging with time. Furthermore we assume that for each $\theta$ fixed, the dynamics $(X_\theta(t))_{t\geq 0}$ is ergodic and therefore (see (18)) we have

$$\frac{1}{T}\int_0^T X_\theta(t)dt \longrightarrow \mathbb{E}\left[X_\theta^*|\theta\right] \qquad \text{as } T\to\infty \tag{35}$$

$$\frac{1}{T}\int_0^T \left(X_\theta(t)\right)^2 dt \longrightarrow \mathbb{E}\left[\left(X_\theta^*\right)^2|\theta\right] \qquad \text{as } T\to\infty \tag{36}$$

where $X_\theta^*$ is the copy number of the measured species at the stationary probability distribution (or equivalently at a sufficiently large time point).

The total cell-to-cell variability at the stationary probability distribution is $\text{Var}(X_\theta^*)$. The law of total variance stipulates that

$$\text{Var}(X_\theta^*) = \text{Var}\left(\mathbb{E}\left[X_\theta^*|\theta\right]\right) + \mathbb{E}\left[\text{Var}(X_\theta^*|\theta)\right] \tag{37}$$

where the quantity $\text{Var}(X_\theta^*|\theta)$ is the conditional variance defined by

$$\text{Var}(X_\theta^*|\theta) = \mathbb{E}\left[(X_\theta^*)^2|\theta\right] - \mathbb{E}\left[X_\theta^*|\theta\right]^2$$

and it measures the variation in $X_\theta^*$ when the parameter $\theta$ is fixed. Hence the only contribution to $\text{Var}(X_\theta^*|\theta)$ comes from sources other than $\theta$, and in this setup the only such source is the *intrinsic noise* introduced by the random timing of reactions. Therefore we term the second term $\mathbb{E}\left[\text{Var}(X_\theta^*|\theta)\right]$ on the r.h.s of Eq. [37] as the aggregate/bulk/population-level measure of the total intrinsic noise in the system.

On the other hand, the first term $\text{Var}\left(\mathbb{E}\left[X_\theta^*|\theta\right]\right)$ on the r.h.s of Eq. [37] measures all the noise due to the randomness in the parameters $\theta$. This is because when we take the conditional expectation $\mathbb{E}\left[X_\theta^*|\theta\right]$ we are averaging-out noise from all other sources except the parameters $\theta$. Due to this reason we term $\text{Var}\left(\mathbb{E}\left[X_\theta^*|\theta\right]\right)$ as the total extrinsic noise in the system. In summary we have the following definition of intrinsic and extrinsic noise

$$\text{Total Extrinsic Noise} = \text{Var}\left(\mathbb{E}\left[X_\theta^*|\theta\right]\right)$$
$$\text{Total Intrinsic Noise} = \mathbb{E}\left[\text{Var}(X_\theta^*|\theta)\right]$$

Of course both these terms add up to the full cell-to-cell variability $\text{Var}(X_\theta^*)$ due to Eq. [37].

The straightforward computation of intrinsic and extrinsic noise is challenging because the experimental data cannot provide a cell population with the same parameters $\theta$ for estimating $\mathbb{E}\left[X_\theta^*|\theta\right]$ and $\text{Var}(X_\theta^*|\theta)$. Fortunately, ergodicity provides us a solution for computing these conditional mean and variance from single-cell time-course data. Specifically, Eq. [35] and Eq. [36] suggest that these conditional mean and variance can be approximated by

$$\mathbb{E}\left[X_\theta^*|\theta\right] \approx \frac{1}{T}\int_0^T X_\theta(t)dt \qquad \text{and} \qquad \text{Var}(X_\theta^*|\theta) \approx \frac{1}{T}\int_0^T \left(X_\theta(t)\right)^2 dt - \left(\frac{1}{T}\int_0^T X_\theta(t)dt\right)^2$$

for large time $T$. Recall that the dynamics of $X_\theta(t)$ is measured in a microscope platform. Therefore, when the measurement noise is negligible, the time integrals above can be straightforwardly computed by the measured single-cell time-course data of $X_\theta(t)$. Finally, the intrinsic noise and extrinsic noise can be evaluated by

$$\text{Total Intrinsic Noise} \approx \mathbb{E}\left(\frac{1}{T}\int_0^T \left(X_\theta(t)\right)^2 dt - \left(\frac{1}{T}\int_0^T X_\theta(t)dt\right)^2\right)$$

$$\text{Total Extrinsic Noise} \approx \text{Var}\left[\frac{1}{T}\int_0^T X_\theta(t)dt\right]$$

for large time $T$.

To further validate this noise decomposition method, let us outline a couple of edge cases where one of the two noise components is zero. First, let us consider the scenario that the dynamics $\{X_\theta(t)\}_{t\geq 0}$ is deterministic, and described by a system of ordinary differential equations (ODEs) that depend on the cell-specific parameter $\theta$. In this case, ergodicity is tantamount to this ODE system having a globally attracting $\theta$-dependent fixed point $\overline{X}_\theta$. One can easily check that in this scenario

$$\lim_{T\to\infty}\frac{1}{T}\int_0^T X_\theta(t)dt = \overline{X}_\theta \qquad \text{and} \qquad \lim_{T\to\infty}\frac{1}{T}\int_0^T \left(X_\theta(t)\right)^2 dt = \overline{X}_\theta^2$$

which shows that the intrinsic noise component would be close to zero (for large $T$). This is what we would expect as the dynamics is deterministic and hence there is no noise due to the random firing of reactions. Now consider another scenario where the parameter is not random, but a deterministic constant $\theta = \theta_c$ which is the same for each cell. In this case the conditional expectation in Eq. [35] becomes an unconditional expectation which computes to a deterministic constant (rather than a random variable). Hence its variance is zero which shows that the extrinsic noise component would be close to zero (for large $T$). Again this is consistent with our expectation because there is no variability in $\theta$.

### References

1. DF Anderson, TG Kurtz, *Stochastic analysis of biochemical systems.* (Springer) Vol. 674, (2015).
2. D Crisan, Particle filters—a theoretical perspective in *Sequential Monte Carlo methods in practice.* (Springer), pp. 17–41 (2001).
3. A Doucet, AM Johansen, A tutorial on particle filtering and smoothing: Fifteen years later. *Handb. nonlinear filtering* **12**, 3 (2009).
4. N Chopin, , et al., Central limit theorem for sequential monte carlo methods and its application to bayesian inference. *The Annals Stat.* **32**, 2385–2411 (2004).
5. A Bain, D Crisan, *Fundamentals of stochastic filtering.* (Springer Science & Business Media) Vol. 60, (2008).
6. M Rathinam, M Yu, State and parameter estimation from exact partial state observation in stochastic reaction networks. *The J. Chem. Phys.* **154**, 034103 (2021).
7. L Duso, C Zechner, Selected-node stochastic simulation algorithm. *The J. chemical physics* **148**, 164108 (2018).
8. ES D'Ambrosio, Z Fang, A Gupta, M Khammash, Filtered finite state projection method for the analysis and estimation of stochastic biochemical reaction networks. *bioRxiv* (2022).
9. J Liu, M West, Combined parameter and state estimation in simulation-based filtering in *Sequential Monte Carlo methods in practice.* (Springer), pp. 197–223 (2001).
10. C Berzuini, W Gilks, Resample-move filtering with cross-model jumps. *Seq. Monte Carlo Methods Pract.* pp. 117–138 (2001).
11. WR Gilks, C Berzuini, Following a moving target—monte carlo inference for dynamic bayesian models. *J. Royal Stat. Soc. Ser. B (Statistical Methodol.* **63**, 127–146 (2001).
12. N Oudjane, C Musso, Progressive correction for regularized particle filters in *Proceedings of the Third International Conference on Information Fusion.* (IEEE), Vol. 2, pp. THB2–10 (2000).
13. Z Fang, A Gupta, M Khammash, Convergence of regularized particle filters for stochastic reaction networks. *SIAM J. on Numer. Analysis* **61**, 399–430 (2023).
14. N Chopin, PE Jacob, O Papaspiliopoulos, Smc2: an efficient algorithm for sequential analysis of state space models. *J. Royal Stat. Soc. Ser. B (Statistical Methodol.* **75**, 397–426 (2013).
15. D CRISAN, J MÍGUEZ, Nested particle filters for online parameter estimation in discrete-time state-space markov models. *Bernoulli* **24**, 3039–3086 (2018).
16. MB Elowitz, S Leibler, A synthetic oscillatory network of transcriptional regulators. *Nature* **403**, 335–338 (2000).
17. A Georgoulas, J Hillston, G Sanguinetti, Unbiased bayesian inference for population markov jump processes via random truncations. *Stat. computing* **27**, 991–1002 (2017).
18. A Gupta, C Briat, M Khammash, A scalable computational framework for establishing long-term behavior of stochastic reaction networks. *PLoS computational biology* **10**, e1003669 (2014).
